# A sequence–structure dual-match logic enables target recognition fidelity by an IS110 transposon

**DOI:** 10.64898/2026.03.19.712850

**Authors:** Xue-Mei Na, Bai-Tao Chen, Han Liu, Yi-Tong Lin, Tai-Jiang Chen, Yuan Yuan, Yu-Hang Zhang, Nan Yang, Yi-Na Xiong, Fei Zhu, Yingying Zhang, Ximin Chi, Zhiyou Zong, Chen Ling

**Affiliations:** State Key Laboratory of Cellular Stress Biology, School of Life Sciences, Xiamen University, Xiamen, Fujian 361102, China; National Engineering Research Center of Industrial Enzymes, Tianjin Institute of Industrial Biotechnology, Chinese Academy of Sciences, Tianjin, China; School of Advanced Interdisciplinary Biomedical Sciences, Xiamen University, Xiamen, Fujian 361102, China

**Author notes:** These authors contributed equally (Xue-Mei Na, Bai-Tao Chen, Han Liu).

## Abstract

IS110 elements employ bridge RNA to direct target site recognition, yet whether additional layers of specificity exist beyond RNA–DNA base pairing is unknown. Here we identified ISPpu10, an IS110 element in *Pseudomonas putida* that is associated with >100-kb segmental amplifications during adaptive laboratory evolution. All insertions mapped to multiple identical copies of a single conserved intergenic sequence harboring a GC-rich 5′ extension that forms a stem-loop. Systematic mismatch and compensatory substitution scanning revealed a sequence**–**structure dual-match logic: base pairing between the bridge RNA and target DNA serves as an RNA-guided match, whereas the target-encoded stem-loop provides an additional specificity determinant. Transplanting this stem-loop into *Escherichia coli* targets improved on-target integration and reduced genome-wide off-target insertion, establishing the stem-loop as a portable structural gate. These findings reveal a structural recognition layer beyond base-pairing, suggesting how bacteria reconcile transposition with genome integrity and offering a strategy for precise genome editing.

## INTRODUCTION

The targeted integration of large, gene-sized DNA payloads remains a critical bottleneck in synthetic biology and therapeutic gene editing.^1^ Although CRISPR-Cas technologies have advanced rapidly toward this goal, including nuclease-mediated endogenous homology-directed repair (HDR), prime editing-assisted site-specific integrase gene editing (PASSIGE), and CRISPR-associated transposases (CASTs), these approaches remain constrained by several limitations: the HDR pathway is largely inactive in postmitotic cells, and both PASSIGE and CASTs leave scars on the genome and require the coordinated action of two independent enzymes, demanding large coding sizes.^2–4^

To this end, a system that combines RNA-guided programmability with the capacity for fragment insertion would offer a promising solution. Recently, the IS110 family of bacterial insertion sequence (IS) elements has been identified and developed into an RNA-guided, reprogrammable genome engineering toolkit. The guiding non-coding RNA, termed bridge RNA (bRNA), possesses two DNA-binding loops, a target-binding loop (TBL) and a donor-binding loop (DBL), each of which pairs with a cognate DNA substrate. The IS110-family transposases have also been termed recombinases, as they mediate recombination between donor DNA (dDNA) and target DNA (tDNA) through two-strand exchange via a Holliday junction intermediate,^5,6^ although an alternative single-strand transfer mechanism has also been proposed.^7^

Recent pioneering studies have shown that IS110 systems bypass the double-strand break (DSB) dependency of CRISPR technologies and mediate large-fragment genomic insertions: IS621 has been validated in bacteria,^5^ whereas ISCro4 does so in human cells.^8,9^ However, a fundamental paradox remains. The bRNA–tDNA pairing interfaces of these systems are short, comprising only 12 bp and 16 bp for IS621 and ISCro4, respectively. For IS621, mismatches occur even within the designated pairing regions,^5^ further underscoring the instability of this interface. Moreover, the stringency of this pairing has not been fully established in currently characterized systems, raising the possibility that the interface may tolerate more mismatches than it appears. From the perspective of bacterial fitness, active transposition of these elements risks disrupting gene function, and a short pairing interface alone seems insufficient to safeguard genome integrity. This led us to ask whether an additional layer of recognition exists beyond bRNA–tDNA sequence pairing.

Addressing this question requires looking beyond the bRNA itself. Previous work on IS110-family elements has been largely RNA-centric, starting from the bridge RNA sequence to predict its DNA partners and thereby emphasizing only those target features explained by direct base pairing.^5,8^ By contrast, little attention has been given to working backward from natural insertion sites and asking what other features they share. This is particularly relevant for IS621 and ISPpu10, both of which have long been known to insert into repetitive extragenic palindromic (REP) sequences.^10,11^ REP elements fold into imperfect palindromes, yet this structural feature is invisible to base-pairing predictions and has therefore been largely overlooked in modular designs.^5^ Systematically analyzing natural IS110 insertion sites beyond base-pairing motifs may thus uncover previously unappreciated determinants of target selection and inform the development of more effective bRNA-guided genome editing tools.

Here, we identify ISPpu10, an IS110 element whose insertions mark the boundaries of two segmental amplifications (227.6 kb and 120.2 kb) in *Pseudomonas putida* KT2440 (hereafter *P. putida*) during adaptive laboratory evolution. We first show that ISPpu10 encodes an autonomous and *trans*-acting IS110-family recombinase that functions across species. Systematic scanning of the bRNA–tDNA interface reveals that base pairing is essential but stringency is asymmetric: the right guide requires the wild-type sequence at nearly every position, while the left guide is permissive except near the recombination site and reprogrammable if base pairing is maintained. Finally, we show that the GC-rich extension of the target REP folds into a stem-loop that acts as a structural gate, enhancing on-target integration and reducing off-target insertion.

## RESULTS

### ISPpu10 insertions mark the boundaries of large-scale segmental amplifications

Our earlier ALE of *P. putida* on xylose as the carbon source yielded a strain carrying a 227.6-kb genomic amplification, detected as an approximately twofold increase in Illumina sequencing coverage across loci PP_5050 to PP_5242.^12^ The amplification boundaries coincided with PP_5050, a 1.3-kb IS annotated as ISPpu10.^11^ However, because ISPpu10 is longer than the Illumina read length, the short reads could not span this element to reveal what the amplified region was joined to, leaving the structural topology unresolved (**Fig. 1A**, top).

**Figure 1.**
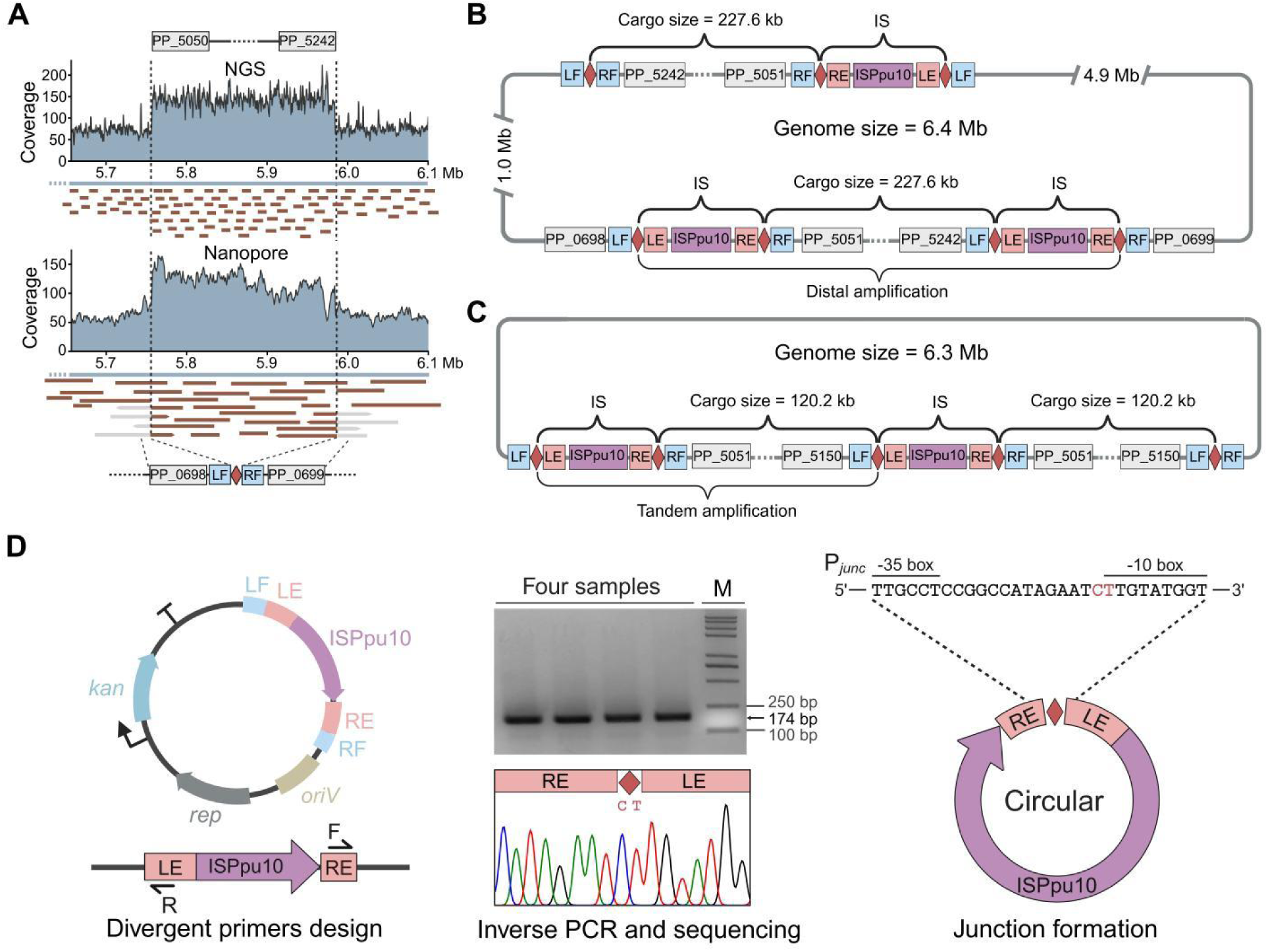
ISPpu10 insertions mark the boundaries of large-scale segmental amplifications. **A.** Comparative genomic coverage of the evolved *P. putida* isolate. Top: Illumina short-read sequencing, re-analyzed from earlier ALE experiments, reveals a stepwise increase in sequencing coverage (227.6 kb) indicative of a segmental amplification but cannot span the repetitive ISPpu10 (PP_5050) boundaries.^12^ Bottom: Oxford Nanopore long-read sequencing bridges the repetitive junctions, resolving the structural organization of the amplification. **B.** Structural map of the 227.6-kb distal segmental amplification. The amplified segment (PP_5050–PP_5242) is flanked by two ISPpu10 copies, in which a de novo ISPpu10 insertion into a downstream REP sequence at PP_5242 defines the second boundary, forming a composite transposon-like architecture (ISPpu10–Cargo–ISPpu10). This unit was mobilized to a distal target locus >1 Mb from the donor site. **C.** Structural map of the 120.2-kb tandem amplification identified in a distinct evolved isolate. The rearrangement manifests as a head-to-tail array, similarly defined by an ISPpu10 insertion into a downstream REP sequence. **D.** Detection of the ISPpu10 circular intermediate. PCR with outward-facing primers (arrows) amplifies the LE–RE junction, which reconstitutes the bRNA promoter P*_junc_* and confirms the excision step. Diamond shapes represent the conserved CT core sequences, the catalytic center of the IS110 recombinase.

Our later Nanopore long-read sequencing successfully bridged these boundaries (**Fig. 1A**, bottom), resolving a distal segmental amplification integrated >1 Mb away. Notably, we detected another ISPpu10 insertion at the 3′ boundary of PP_5242, precisely within a REP sequence (**Fig. 1B**). Upon integration, the element is flanked by the two halves of the genomic REP target, which we term the left and right flanks (LF and RF), adjacent to the element’s left and right ends (LE and RE). This generates an architecture resembling a composite transposon, which we term ISPpu10–Cargo–ISPpu10 (**Fig. 1B**). In a separate ALE experiment, a distinct evolved isolate exhibited a 120.2-kb tandem amplification, manifesting as a head-to-tail ISPpu10–Cargo–ISPpu10 array similarly defined by a novel ISPpu10 insertion into a downstream REP site (**Fig. 1C**). In both cases, the rearrangements were flanked by ISPpu10 insertions into specific REP sequences, consistent with its previously reported target specificity.^11^

IS621 has been reported to catalyze the excision process through low basal expression of the recombinase and bRNA.^13^ To test whether the native ISPpu10 element catalyzes excision, we cloned it, encompassing its LE and RE together with the genomic LF and RF flanks, into a medium-copy pBBR1 plasmid and introduced the construct into *Escherichia coli* (*E. coli*) TOP10, which lacks IS110-family elements. PCR amplification and Sanger sequencing confirmed the precise fusion of LE and RE, indicative of a circular excision intermediate (**Fig. 1D**). Although circularization is employed by other mobile elements including IS3 to transiently generate hybrid promoters,^14^ this step is obligatory in the IS110 life cycle: the precise LE–RE fusion assembles the complete donor DNA substrate for recombination, and the resulting junction promoter (P*_junc_*) drives transcription of the bridge RNA required for sequence recognition.^5,13,15^

Thus, ISPpu10 encodes a functional IS110-family recombinase capable of precise excision through a circular intermediate.

### ISPpu10 is an autonomous and *trans*-acting recombinase

To characterize the transposition activity of ISPpu10, we first defined the boundaries of its functional elements. The target substrate is the genomic REP sequence,^11^ whereas the donor substrate is the circular intermediate formed by LE–RE fusion; based on a promoter predicted to span the core dinucleotide (CT) on this intermediate, we defined the bRNA transcription unit as extending from the P*_junc_*+1 site to the start of the CDS (**Fig. 2A**). To identify the base-pairing regions between the bRNA and the donor and target DNA substrates, we mapped the bRNA onto these two substrates following the pairing logic established for IS621.^5^ This mapping defined the left target (LT) and right target (RT) within the REP sequence, the left donor (LD) and right donor (RD) at the termini of LE and RE. It also revealed two base-pairing loops in the bRNA: within the target-binding loop (TBL), the left target guide (LTG) and right target guide (RTG) pair with LT and RT on opposite strands of the tDNA, whereas within the donor-binding loop (DBL), the left donor guide (LDG) and right donor guide (RDG) pair with LD and RD on opposite strands of the dDNA (**Fig. 2A, B**). The interfaces tolerate specific deviations from canonical Watson–Crick pairing: the LTG–LT interface accommodates a U·G wobble, mirroring the G·T wobble found at the corresponding interface of IS621, whereas the RDG–RD interface uniquely contains a G·G mismatch.^5^

**Figure 2.**
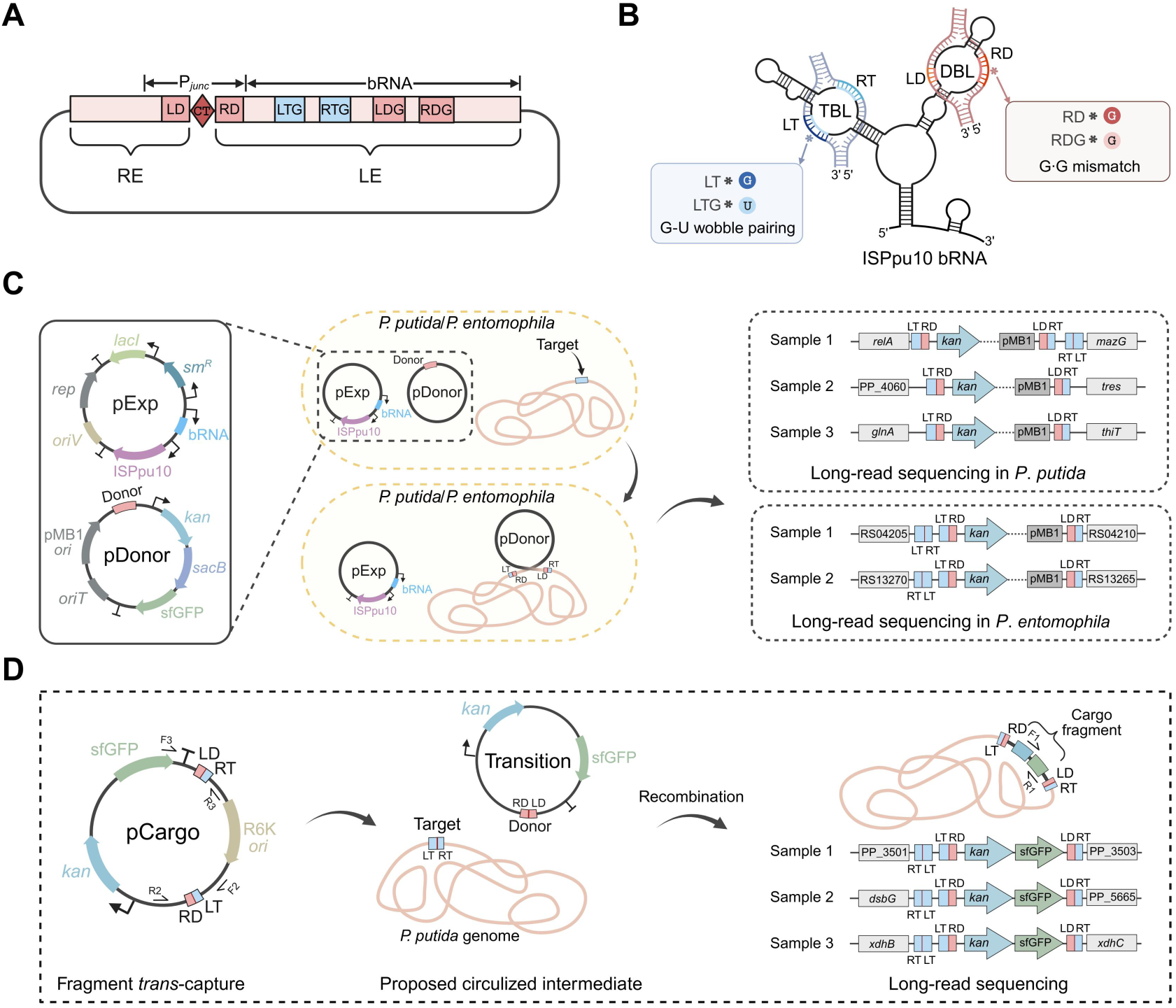
ISPpu10 is an autonomous and *trans*-acting recombinase. **A.** Schematic of the excised circle showing the fused LE–RE junction. The reconstituted hybrid promoter P*_junc_* (−35 box from RE, −10 box from LE) drives transcription of the bRNA from the +1 site through LE to the start of the CDS. CT, central conserved core. **B.** Architecture of the ISPpu10 bRNA and its interaction with donor and target DNA. The bRNA contains a TBL and a DBL. The TBL base-pairs with the LT and RT on opposite DNA strands; the DBL base-pairs with the LD and RD. The interfaces tolerate specific non-canonical pairs, including a G·U wobble at the LT interface and a G·G mismatch at the RD interface. **C.** Site-specific genomic integration in *P. putida*. Co-expression of ISPpu10 and its bRNA with a suicide donor plasmid carrying an antibiotic resistance marker results in integration into a chromosomal REP target. Colony PCR confirmed integration in 87.5% of the recovered colonies, and long-read sequencing of three randomly selected colonies verified precise donor insertion. Inset: heterologous expression in *Pseudomonas entomophila*, which harbors identical REP sequences but lacks ISPpu10, also yielded correct genomic insertions, demonstrating that the recombinase and its bRNA are sufficient without factors specific to *P. putida*. Full gel images are provided in Fig. S2. **D.** Fragment *trans*-capture and integration from a split-donor substrate. A suicide plasmid carrying a split-donor cassette (LE–cargo–RE), in which LE and RE flank a cargo sequence rather than a complete IS element, was introduced into *P. putida* by electroporation. Three colonies carrying both expected integration junctions were identified by junction-spanning PCR, and long-read sequencing confirmed precise backbone-free insertion of the cargo into a chromosomal REP target.

To determine whether ISPpu10 functions as an autonomous recombinase, we co-expressed ISPpu10 and its bRNA from a helper plasmid (pExp) together with a suicide donor plasmid (pDonor) carrying an antibiotic resistance marker in *P. putida*. Because pDonor cannot replicate autonomously, cells could be recovered on antibiotic selection only if pDonor had integrated into the chromosome. Integration into a chromosomal REP target was detected in 87.5% of the recovered colonies by colony PCR, and long-read sequencing of three randomly selected colonies confirmed precise donor insertion (**Fig. 2C**). We next expressed the same two-plasmid system in *Pseudomonas entomophila*, a related species that harbors identical REP sequences but lacks ISPpu10; correct insertions were likewise obtained in this heterologous host (**Fig. 2C**), demonstrating that the recombinase and its bRNA are sufficient for integration without requiring factors specific to *P. putida*.

Finally, the composite ISPpu10–Cargo–ISPpu10 architecture described above (**Fig. 1B**) suggests that the mobilized donor might involve recognition ends that are separated by the cargo rather than adjacent within a single element. To test whether ISPpu10 can act on such a donor *in trans*, we electroporated into *P. putida* a suicide plasmid carrying a split-donor cassette (LE–cargo–RE), in which LE and RE flank a cargo sequence, together with a plasmid expressing ISPpu10 and its bRNA. Three colonies carrying both the LT–RD and LD–RT junctions were identified by junction-specific PCR screening (**Fig. S1**), and long-read sequencing confirmed backbone-free integration of the cargo into chromosomal REP sequences (**Fig. 2D**). Notably, in all three colonies, the integration site lay within 100 bp of a second intact REP sequence (58 bp, 58 bp, and 24 bp, respectively), although the basis for this proximity remains unclear. This result demonstrates that ISPpu10 possesses *trans* activity: it can capture a LE–cargo–RE fragment and integrate it into the genome without requiring the donor ends to be present *in cis* on a complete IS element.

### The target stem-loop is conserved across ISPpu10 homologs

To characterize the genomic context of ISPpu10 integration, we mapped all REP target sites across the *P. putida* genome and found that the target sequences are distributed throughout the genome (**Fig. 3A**). Consistent with the intergenic location of REP sequences, 97.1% of ISPpu10 targets reside outside coding regions, and 40.8% occur as convergent pairs spaced less than 200-bp apart (**Fig. 3B**), an arrangement that may increase the local density of integration sites.

**Figure 3.**
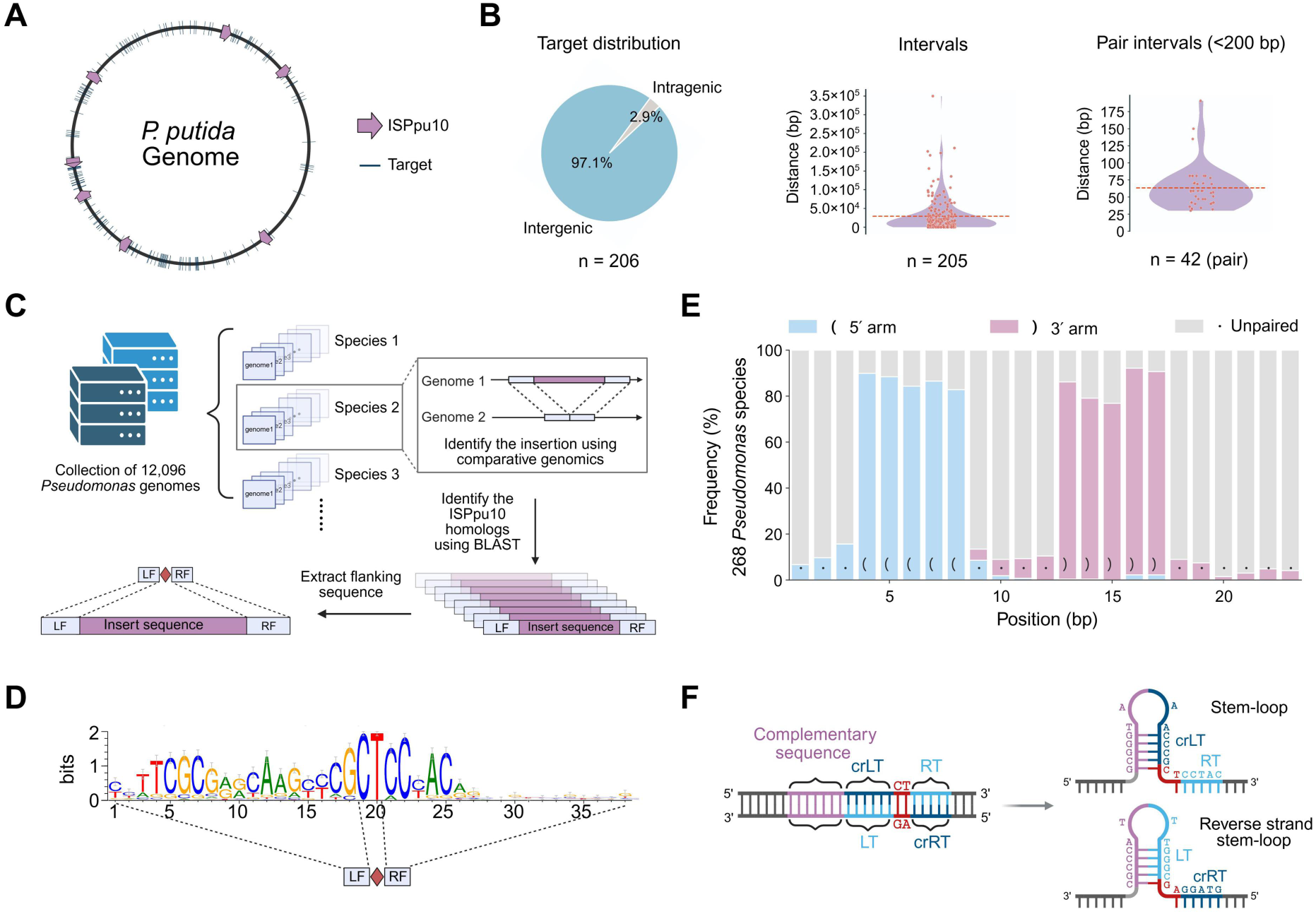
Conserved target-site stem-loop architecture in *Pseudomonas* genomes. **A.** Genome-wide distribution of ISPpu10 REP target sites in *P. putida*. **B.** Genomic context of predicted REP target sites in *P. putida*. Among all predicted REP target sites genome-wide, 97.1% reside outside coding regions, and 40.8% occur as convergent pairs spaced less than 200 bp apart within intergenic regions. **C.** Workflow for the systematic identification of ISPpu10 homologs. A dataset comprising 12,096 *Pseudomonas* genomes was curated and grouped into species-specific clusters. Comparative genomics within these clusters (exemplified by Species 2) enabled the detection of sequence insertions, and ISPpu10 homologs were subsequently identified using BLAST. This pipeline allowed for the extraction and alignment of the insertion sequences alongside their adjacent left and right flanking regions (LF and RF). **D.** Sequence logo showing the conservation of target sequences. REP target sequences were reconstructed for 268 homologous ISPpu10 elements from related *Pseudomonas* species by joining the sequences flanking each insertion site. **E.** Conserved secondary structure at the target sites of homologous ISPpu10 elements. The frequency of each predicted secondary-structure state at each position (23-bp window) is shown using dot-bracket notation: blue, 5′ stem; pink, 3′ stem; grey, unpaired. The consensus structure reveals a highly conserved stem-loop characterized by a ∼5-bp base-paired stem (positions 4–8 and 13–17) and a predominantly unpaired loop (positions 9–12). Structural conservation (84.63% covariation among paired positions) exceeded sequence conservation (63.10% average pairwise identity) across the same region. **F.** Structural model of the target-site architecture. Intra-strand complementarity drives the folding of a conserved 5′ stem-loop within the target DNA.

Although REP sequences have long been recognized as IS110 targets,^10,11^ the structural features of the REP target, beyond its base pairing with the bRNA, have received little attention. We therefore sought to characterize ISPpu10 target sequences in an unbiased manner. Comparative genomics were performed across 12,096 *Pseudomonas* genomes in NCBI, resolving insertion events as genomic segments present in some genomes but absent from others (**Fig. 3C**). Matching these insertion events to the ISPpu10 recombinase sequence by BLAST identified 268 homologous elements, for each of which the insertion boundaries could be defined. For each element, we extracted 20 bp of host sequence from each side of the insertion, removed one copy of the duplicated CT dinucleotide (the 2-bp target-site duplication), and joined the two flanks to reconstruct a 38-bp target sequence (**Fig. 3D**). Multiple sequence alignment of these 268 sequences revealed, in addition to the left target (LT) and right target (RT) regions, a conserved 5′ sequence whose function was unclear. We therefore focused our structural analysis on the 23-bp conserved core of the alignment, which for ISPpu10 corresponds to its annotated target in ISfinder.^16^ Strikingly, structural conservation across the 268 sequences exceeded sequence conservation: the covariation rate among paired positions was 84.63%, whereas the average pairwise identity of the same region was only 63.10% (**Fig. 3E**). The marked excess of structural over sequence conservation indicates that the stem-loop fold is under stronger selective constraint than its primary sequence. Consistent with this, in the ISPpu10 target itself, the conserved 5 ′ sequence is complementary to crLT (the reverse complement of LT, hereafter crLT), and intra-strand base pairing between them drives folding of a 5′ stem-loop (**Fig. 3F**).

Collectively, ISPpu10 targets are characterized by a conserved 5′ stem-loop. This raises the question of whether the target-encoded stem-loop contributes to target recognition beyond base pairing.

### Base pairing and the target stem-loop act as complementary recognition determinants

To test whether the target-encoded stem-loop contributes to ISPpu10 function, we developed a dual-color reporter in which recombination switches fluorescence from sfGFP to mCherry: recombination excises the donor from a permissive loop of sfGFP, abolishing green fluorescence, while promoter translocation activates a promoter-less mCherry (**Fig. 4A**, **Fig. S3–7**). Replacing the GC-rich stem-forming region of the target with a poly(A) sequence did not affect the LTG–LT base pairing but disrupted the intra-strand pairing between crLT and its 5′ complementary sequence that forms the stem (**Fig. 4B**). Disrupting the stem-loop in this way significantly reduced integration efficiency (**Fig. 4B**), indicating that the stem-loop promotes recombination.

**Figure 4.**
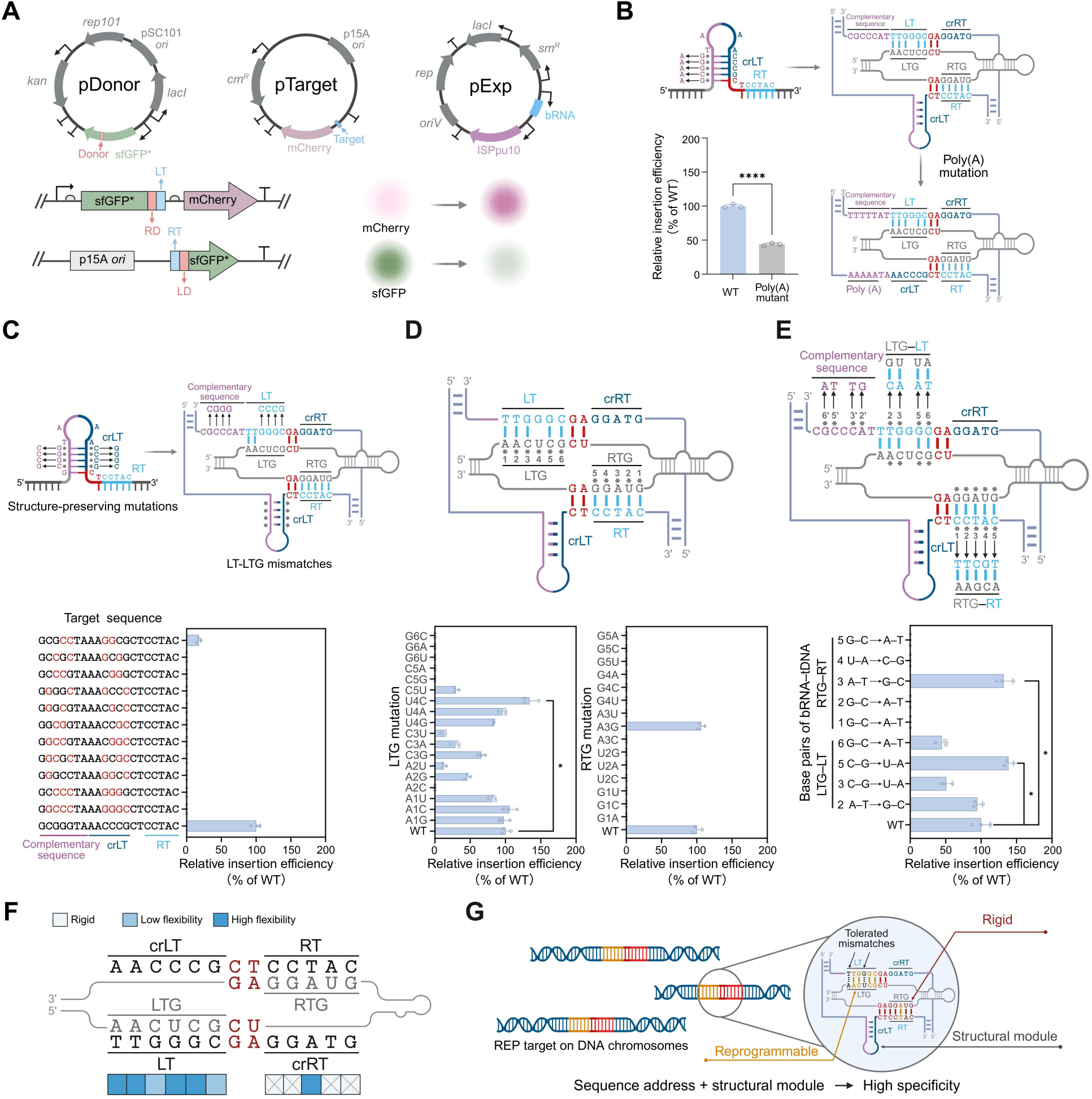
Sequence and structural determinants of ISPpu10 target recognition. **A.** Dual-color reporter assay. Recombination between the donor (pDonor) and target (pTarget) plasmids excises the donor from the sfGFP CDS and translocates a promoter to activate mCherry, converting green to red fluorescence. RD–LT and RT–LD junctions indicate recombination. **B.** The target-encoded stem-loop enhances integration efficiency. Replacing the GC-rich stem-forming region with a poly(A) sequence, which abolishes stem-loop formation, reduced integration to 43.85% of the wild-type level. **C.** Stem base-pair inversion preserves stem-loop folding while disrupting bRNA–target pairing, severely compromising activity. Red nucleotides indicate altered positions. **D.** Mismatch tolerance across the bRNA–target interface. Single-base substitutions show that the LTG tolerates mismatches at distal positions but is constrained near the CT core, whereas the RTG is stringent except at position 3. **E.** Base-pair exchange with complementarity maintained. The LTG–LT interface tolerates exchange, whereas the RTG–RT interface is sequence-specific. **F.** Flexibility map integrating the mismatch and exchange results. Dark blue, high flexibility; light blue, low flexibility; frown symbol, rigid. The left interface (LT): positions 1 and 4 tolerate all mismatches, positions 2 and 5 tolerate exchange, position 5 favoring a weaker A–U configuration, and positions 3 and 6 retain ∼47.81% activity. The right interface (RT) is rigid except position 3, where a T–A to C–G exchange increases activity to ∼131.97%. **G.** Dual-match logic model. Base pairing defines the genomic address; the 5′ stem-loop acts as a structural module that enhances specificity. Data in b, d and e are mean ± s.d. (n = 3); \**p* < 0.05, \*\*\*\**p* < 0.0001 (two-tailed Student’s *t*-test).

To ask whether the stem-loop alone can sustain activity when base pairing is weakened, we inverted the base pairs of the stem in successive steps (two, three, or four pairs). These manipulations preserved stem-loop folding while disrupting the LTG–LT base pairing at the inverted positions (inversion variants marked with asterisks). Of eleven inversion variants, ten abolished recombination; the sole exception, which inverted the two pairs nearest the loop, retained 17.58% activity, consistent with a residual wobble pairing at one inverted position (**Fig. 4C**). These results show that the stem-loop cannot compensate for the loss of key base pairing.

Together, these experiments define a functional hierarchy: base pairing is required for recombination, whereas the stem-loop enhances recombination efficiency once the correct target is engaged.

To map the base-pairing constraints, we performed single-base saturation mutagenesis across the bRNA TBL against an unmodified wild-type target that retains the 5′ stem-loop, thereby isolating base pairing as the sole experimental variable. This scan revealed a marked asymmetry between the two interfaces. The LTG exhibited a position-dependent tolerance for mismatches: the distal positions 1 and 4 fully tolerated all substitutions, positions 2 and 3 accepted multiple mismatches with reduced activity, and position 5 retained activity only upon a C-to-U substitution, which forms a G·U wobble, whereas position 6, adjacent to the CT core, was strictly intolerant of any mismatch (**Fig. 4D**). By contrast, the RTG required base pairing at nearly every position, with the sole exception of an A-to-G substitution at position 3, which forms a G·T wobble and supports insertion with comparable efficiency (**Fig. 4D**).

We next asked whether the two interfaces can be reprogrammed while maintaining guide–target base pairing. We exchanged base pairs at the LTG–LT and RTG–RT interfaces under two rules: complementarity was preserved, while the bases were switched to maximally different nucleotides, so that a C–G pair became a U–A pair and an A–T pair became a G–C pair. We tested RTG positions 1 to 5, and LTG positions 2, 3, 5, and 6, excluding positions 1 and 4 because they already tolerate mismatches. At the LTG–LT interface, every exchanged position retained activity, with positions 3 and 6 reduced to roughly 47.81% of the wild-type level, whereas the C–G to U–A exchange at position 5 increased activity to about 138.58% of wild type (**Fig. 4E**). At the RTG–RT interface, the A–T to G–C exchange at position 3 likewise increased activity to about 131.97% of wild type, whereas exchanges at the other four positions abolished activity (**Fig. 4E**). Together with the mismatch scan, these exchange results define a flexibility map of the two interfaces (**Fig. 4F**): both interfaces demand base pairing, whereas the LTG–LT tolerates sequence variation once complementarity is maintained, the RTG–RT additionally requires a specific sequence, the basis for which remains unclear.

Together with the structural contribution of the target-encoded stem-loop (**Fig. 4B, C**), these results establish a dual-match logic for ISPpu10 target selection, in which base pairing defines the target site and the stem-loop enhances recombination efficiency (**Fig. 4G**).

### Modeling suggests the target stem-loop engages the recombinase

To explore how the target-encoded stem-loop might contribute to ISPpu10 function, we performed molecular dynamics (MD) simulations of the recombinase dimer bound to the bRNA and double-strand tDNA, with the 5′ stem-loop folded on the forward strand (**Fig. 5A**). We compared this assembly with two controls: a sequence-length-preserving control in which the complementary sequence that forms the stem was replaced with poly(A), and a truncation control in which this sequence was deleted. In all models, the LT and RT recognition sequences and their base pairing with the bRNA were preserved, so that any difference could be attributed to the stem-loop.

**Figure 5.**
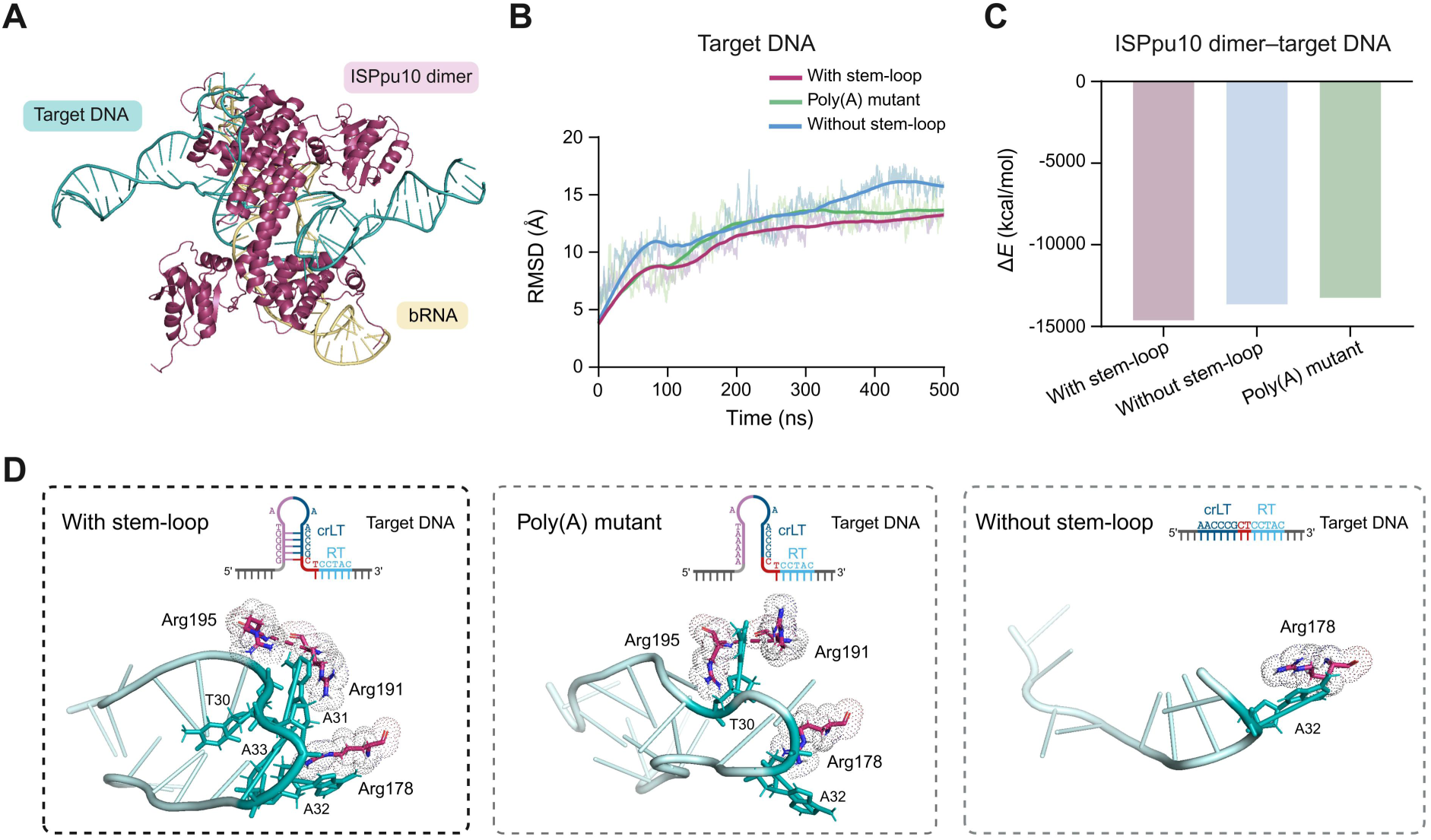
Predicted structural engagement of the target stem-loop by the recombinase. **A.** Simulated dimeric structure of ISPpu10 with bRNA and tDNA. **B.** MD simulations of the ISPpu10–bRNA–tDNA complex. The presence of the 5′ target hairpin reduces the RMSD of the target DNA compared to the hairpin-free and poly(A) mutant systems. **C.** Average protein–target DNA interaction energy calculated from the final 50-ns trajectory. The stem-loop assembly shows stronger interaction energies than the controls. **D.** Structural snapshots from the MD trajectory showing the predicted arginine–loop interactions. In the stem-loop system, the initially designed G29–C34 and G28–C35 pairs undergo a register shift to T30–C34, G29–C35, and G28–C36 hydrogen bonds, so that the hairpin fold is retained; Arg178 and Arg191 form cation–π contacts with A32/A33 and A31, respectively, and Arg195 engages the T30 phosphate. In the poly(A) control, the loop structure is disrupted and the contacts become predominantly electrostatic; the truncation control retains only a single Arg178–A32 contact.

Despite identical base-pairing interfaces, the presence of the stem-loop reduced the root-mean-square deviation (RMSD) of the tDNA relative to both controls, indicating greater rigidity in the simulation (**Fig. 5B**). Interaction energy calculations likewise showed stronger protein–tDNA interactions in the stem-loop assembly (**Fig. 5C**). In the equilibrated trajectory, the stem-loop underwent a register shift: upon contact with the surrounding arginines, the initially designed G29–C34 and G28–C35 base pairs rearranged into T30–C34, G29–C35, and G28–C36 hydrogen bonds, so that the hairpin fold was retained with a shifted register (**Fig. 5D**). The loop residues engaged a specific arginine network, Arg178 and Arg191 forming cation–π contacts with A32/A33 and A31, respectively, and Arg195 engaging the T30 phosphate, whereas the poly(A) control failed to retain the loop structure and its contacts became predominantly electrostatic, and the truncation control retained only a single Arg178–A32 contact. These simulations thus predict that the stem-loop stabilizes the recombinase–DNA complex through its capacity to re-pair and maintain a folded loop conformation that supports a specific arginine interaction network (**Fig. 5D**).

### The stem-loop is a portable safety module that suppresses off-target integration

Naturally occurring stem-loop targets are rare in the *E. coli* genome. We therefore selected four genomic sites that can be targeted by reprogrammed bRNAs based on the constraint rule defined in **Fig. 4**. Using CRISPR-Cas9,^17^ we introduced a sequence complementary to crLT immediately upstream of each site (**Fig. S8–11**), generating four isogenic strain pairs in which a native target (Target 1–4) is compared with its stem-loop-modified counterpart (S-Target 1–4) at each locus (**Fig. 6A**).

**Figure 6.**
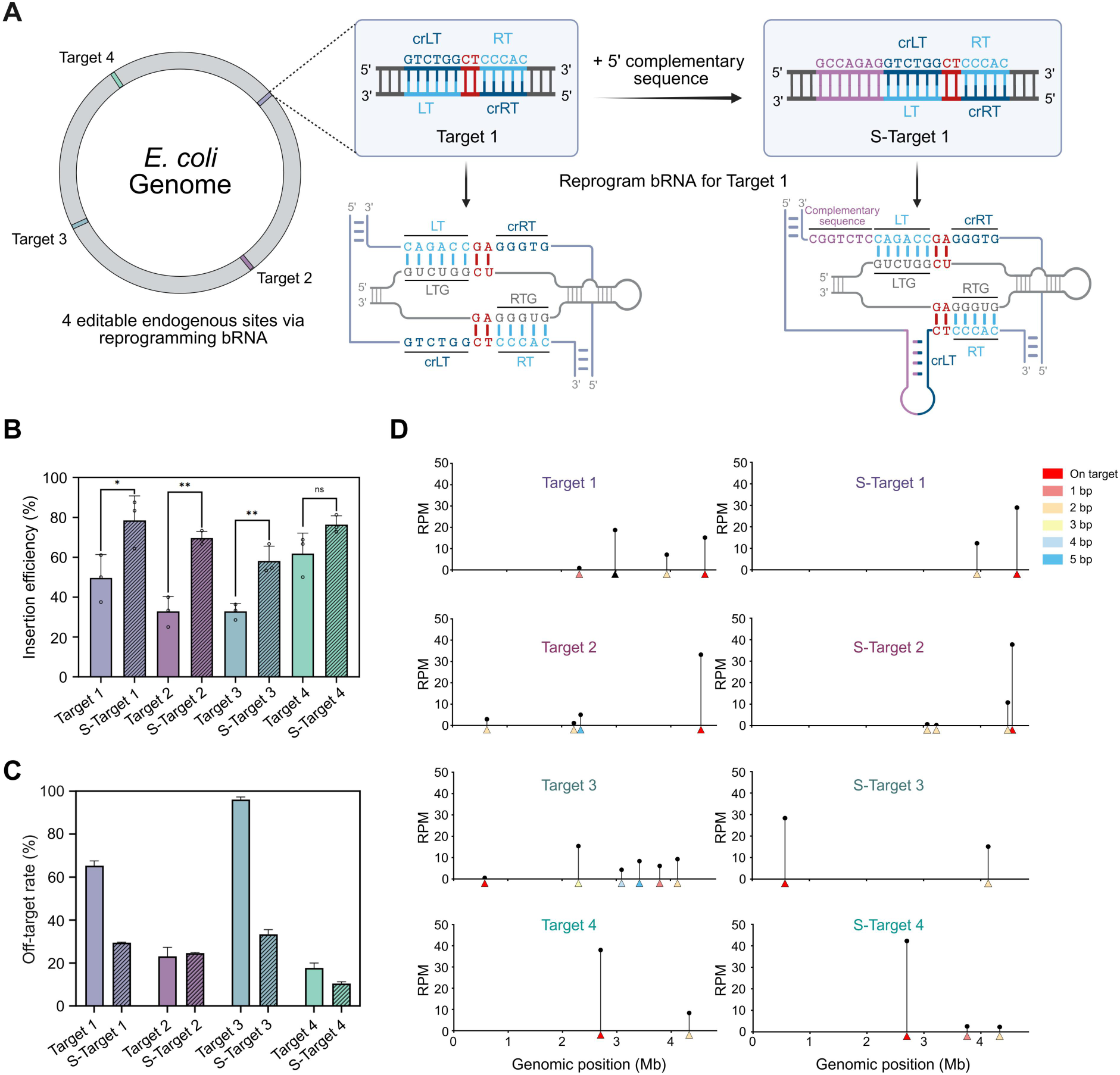
The target-encoded stem-loop is a portable structural gate. **A.** Design of heterologous target sites in the *E. coli* TOP10 genome. Four genomic loci were selected based on sequence complementarity to the ISPpu10 bRNA (Targets 1–4). At each locus, a sequence complementary to crLT was introduced immediately upstream of the bRNA-pairing sequence, thereby reconstituting the 5′ stem-loop and generating four stem-loop-modified targets (S-Targets 1–4). Target 1 and S-Target 1 are shown as an example. **B.** Integration efficiency at native versus stem-loop-modified targets. The presence of the stem-loop increased integration efficiency at all four loci, three of which reached statistical significance. Bars represent mean ± s.d. (n = 3 biological replicates); ns, not significant; \**p* < 0.05, \*\**p* < 0.01 (two-tailed Student’s *t*-test). **C.** Off-target insertion rates determined by deep sequencing. At three of four loci, the stem-loop-modified version exhibited a substantially lower off-target rate than its native counterpart, whereas Target 2 showed a comparable off-target rate. Error bars denote the absolute difference between left and right junction-spanning reads. **D.** Genome-wide off-target profiling by deep sequencing, shown as Manhattan plots across the genome. The y-axis indicates off-target reads per million (RPM), and off-target sites are color-coded by their relationship to the target: on-target and one- to five-base-pair differences are indicated, whereas black marks insertions that are unrelated to the target and independent of the CT core. At all four loci, the stem-loop elevated on-target RPM. Residual off-target insertions were confined to sites carrying one or two mismatches, whereas the more divergent off-target sites, including the unrelated insertions observed in the unmodified target, were largely eliminated.

Across all four loci, the presence of the stem-loop increased integration efficiency relative to the unmodified target, with three of four pairs reaching statistical significance (**Fig. 6B**, **Fig. S8–11**). To assess whether the stem-loop also suppresses off-target insertion, we performed deep sequencing of integration junctions from all eight strains. At three of four loci, the stem-loop-modified target exhibited a substantially lower off-target rate than its native counterpart, whereas Target 2 showed a comparable off-target rate (**Fig. 6C**). Genome-wide analysis further revealed that the stem-loop reshaped the off-target landscape: on-target reads were elevated at all four loci, while off-target insertions were restricted to sequences differing from the target by only one or two base pairs, with the more distant off-target sites observed in the unmodified targets largely eliminated (**Fig. 6D**).

These results demonstrate that the target-encoded stem-loop is a portable specificity module, whose enhancement of on-target integration and suppression of off-target insertion can be reconstituted in a heterologous host by introducing a sequence that pairs with crLT to form the stem-loop. Together with the dual-match logic defined above, this establishes the stem-loop as a structural gate: a modular specificity determinant that operates independently of the REP sequence environment.

## DISCUSSION

The identification of ISPpu10 at the boundaries of two >100-kb segmental amplifications emerged not from computational mining but from a beneficial genomic rearrangement captured during ALE (**Fig. 1**). Although whether ISPpu10 drives or merely accompanies these duplications remains to be determined, the IS–Cargo–IS architecture and our *trans*-capture experiments suggest that ISPpu10 can mediate the underlying rearrangements, through excising large fragments and inserting them into new loci. These observations raise the possibility that ISPpu10 can act on fragments on the order of 100 kb. By comparison, the largest reported insertions by IS110 elements remain below 5 kb,^5,8,9^ and a recent protein-engineered IS621 reached 10 kb only at low efficiency.^18^ We reason that the target-encoded stem-loop may contribute to this large-scale capacity.

The distribution of REP sequences across the *P. putida* genome raises the question of ISPpu10’s biological function. Among the 206 REP sequences, 97.1% reside outside coding regions, with the remaining 2.9% in predicted genes of unknown function. This localization, together with the observation that a beneficial distal amplification was selected under growth pressure during ALE, suggests that ISPpu10 may serve as a mechanism for acquiring or mobilizing functional DNA, thereby contributing to the metabolic versatility of *P. putida*.^19^ Intriguingly, ISPpu10’s association with these amplifications came to light during an ALE: the amplified segment contains *aroB*, and overexpression of this gene relieved the growth pressure by alleviating a metabolic bottleneck.^12^ This observation, although the captured sequence derives from native genes, lends further support to our sequence-capture hypothesis. Notably, several REP-flanked regions exhibit GC content substantially below the genomic average and encode complete metabolic pathways, including fatty acid biosynthesis, the *β*-ketoadipate pathway, and a branched-chain keto acid dehydrogenase complex (**Fig. S12**). The placement of these low-GC metabolic clusters between REP sequences raises the possibility that ISPpu10-mediated *trans*-capture contributed to assembling the metabolic repertoire, although the timing of these events remains unresolved. A second line of support comes from the donor interface: the DBL–dDNA interface is fully reprogrammable at all nine positions, including a naturally occurring G·G mismatch (**Fig. S13**). This flexibility, together with the recent observation that the IS621 DBL alone can mediate both excision and integration in the absence of a functional TBL,^13^ suggests that the system is inherently permissive toward a broad range of DNA substrates.

The target-encoded stem-loop that gates ISPpu10 integration illustrates an economical strategy achieving target specificity without enlarging the element’s coding capacity. Larger transposons such as Tn7,^20^ recruit dedicated accessory proteins for target selection, whereas ISPpu10, a minimal transposon element, achieves an additional layer of specificity by exploiting a structural feature already present in the target DNA itself. Across the target sites of 268 homologous elements, the stem-loop is conserved more strongly at the level of structure than of sequence, suggesting that exploiting nucleic acid structure for integration fidelity may be a recurrent solution in autonomous elements; however, direct functional evidence is currently available only for ISPpu10.

Finally, the target-encoded stem-loop gate offers a conceptual blueprint for enhancing genome-editing specificity. Most programmable nucleases and recombinases rely solely on sequence complementarity for target selection, leaving them vulnerable to off-target activity at partially matched sites, as we observed when targeting the *E. coli* genome without the stem-loop (**Fig. 6C**). ISPpu10 demonstrates that embedding a structural requirement within the target itself, resulting in a sequence-and-structure two-factor authentication, can substantially suppress off-target insertion without sacrificing on-target efficiency (**Fig. 6B–D**). More broadly, the combination of features uncovered here, namely a target-encoded specificity gate, an asymmetric target interface that facilitates reprogramming, a fully flexible donor interface, and the potential for large-scale rearrangements, establishes ISPpu10 as a promising platform for high-fidelity genome engineering.

## METHODS

### Analysis of short-read sequencing coverage

Raw sequencing reads underwent quality control using fastp^21^ to remove low-quality bases and adapter sequences. The filtered paired-end reads were aligned to the *P. putida* reference genome (NC_002947.4) using BWA-MEM^22^ (v0.7.17) with default parameters. Post-alignment processing, including SAM-to-BAM conversion, coordinate sorting, and indexing, was performed using SAMtools^23^ (v1.6). Genome-wide per-base sequencing depth was calculated using the samtools depth command. To generate high-resolution coverage tracks for specific genomic regions, raw depth data were processed using custom Python scripts employing Pandas^24^ (v2.2.2) and Matplotlib (v0.1.7). To mitigate high-frequency noise (such as PCR artifacts) and visualize broad coverage trends, a two-step smoothing strategy was implemented: first, a rolling median filter (window size = 51 bp) was applied to attenuate outliers; subsequently, the data were smoothed using a rolling mean filter (window size = 1 kb). The final coverage profiles were visualized as vector-based area charts (SVG/PDF) to ensure high quality for figure assembly.

### Long-read sequencing and structural variation analysis

Genomic DNA from the first two evolved isolates was sequenced on the Oxford Nanopore platform by Plasmidsaurus. All subsequent long-read sequencing in this study was performed in our lab on the QitanTech QNome-3841 platform. DNA libraries were prepared using the QitanTech Library Preparation Kit following the manufacturer’s instructions. Sequencing was performed on a QCell-3841 flow cell, and basecalling was executed using the QitanTech proprietary basecaller to generate raw FASTQ reads. The quality distribution of raw long-read sequencing data was initially assessed. To maximize read length and continuity—which are critical for resolving complex structural variations—no aggressive quality filtering or trimming was performed, as the mean Q-scores were confirmed to be >10. The raw reads were aligned to the *P. putida* reference genome (NC_002947.4) using BWA-MEM (v0.7.17) with default parameters. Sequencing depth calculation and the identification of high-coverage regions followed the same pipeline as described for the short-read data. To investigate complex structural features, supplementary alignments (SA) were specifically extracted and quantified using custom Python scripts to identify potential chimeric mapping events. Long reads associated with high-coverage SA clusters were subsequently targeted and extracted. The sequences of these specific supplementary alignments were manually inspected to precisely define the insertion boundaries and resolve the sequence architecture of the chromosomal structural variations.

### Detection of circularized intermediates

The native ISPpu10 element, including its flanking LT and RT sequences (LT–LE–ISPpu10–RE–RT), was amplified from *P. putida* genomic DNA. The fragment was cloned into the pBBR1 plasmid backbone via Gibson assembly and transformed into *E. coli* TOP10. Positive colonies were validated by colony PCR and Sanger sequencing (Tsingke). To assay for circularization, sequence-verified *E. coli* TOP10 colonies were cultured in LB medium supplemented with 50 μg mL^-1^ kanamycin at 37℃ for 12 h. Plasmid DNA was extracted and analyzed by PCR using divergent primers designed to amplify the circular junction. Products were resolved on 2% agarose gels, and the specific fusion of the LE and RE was confirmed by Sanger sequencing. Primers and plasmid construction details are described in Supplementary Table 2 and 3.

### RNA secondary structure prediction

The secondary structures of the candidate RNA sequences were predicted using the ViennaRNA Package^25^ (v2.5.0). Specifically, the RNAfold program was employed to calculate the thermodynamically most stable structures based on the minimum free energy (MFE) algorithm. All predictions were performed using default thermodynamic parameters.

### Identification and targeting of endogenous *P. putida* and *P. entomophila* genomic loci

To assess the genomic integration capability of the native ISPpu10 system in *P. putida* and *P. entomophila*, a dual-plasmid system was established. This system consisted of a replicable expression plasmid encoding both the bRNA and ISPpu10, and a non-replicable donor plasmid harboring the donor sequence along with a sfGFP reporter gene. Following plasmid construction, the expression plasmid was directly electroporated into *P. putida* or *P. entomophila* to generate recipient strains. The donor plasmid was introduced into *E. coli* WM3064 via electroporation to serve as the donor strain. Successful transformation was confirmed by colony PCR in all cases. Verified donor strain was cultured overnight in LB supplemented with 0.3 mM diaminopimelic acid (DAP) and 50 µg mL^-1^ kanamycin at 37℃ with shaking at 220 rpm. Verified recipient strains were cultured overnight in LB containing 100 µg mL^-1^ spectinomycin at 30℃ with shaking at 220 rpm. For conjugation, 1.5 mL aliquots of each overnight culture were pelleted by centrifugation at 6,000 rpm for 1 min, supernatants were discarded, and cell pellets were washed with 1 mL PBS. The wash step was repeated once. After the final wash, donor and recipient cells were each resuspended in 100 µL PBS, mixed thoroughly, and spotted onto LB agar plates containing 0.03 mM DAP and 5 µg mL^-1^ kanamycin for 12 h of mating. Following conjugation, the cell biomass was scraped from the mating plates, resuspended in 100 µL PBS containing 10 mM IPTG, and spread onto selection plates (LB agar supplemented with 50 µg mL^-1^ kanamycin and 100 µg mL^-1^ spectinomycin). Plates were incubated at 30℃ for 48 h. The resulting single colonies were examined under blue light for sfGFP fluorescence. sfGFP-positive colonies were subsequently streaked and subjected to colony PCR using primers flanking the donor sequence on the donor plasmid. The absence of a PCR product indicated successful donor excision, whereas a detectable amplicon signified retention of the intact donor plasmid. Colonies yielding no PCR product were inoculated into LB broth containing 50 µg mL^-1^ kanamycin and 100 µg mL^-1^ spectinomycin, cultured overnight, and subjected to genomic DNA extraction followed by third-generation sequencing (Qitan Technology, Chengdu, China). Sequencing reads were analyzed to validate successful donor plasmid integration and to confirm that integration occurred at the predicted wild-type ISPpu10 target sites. Primers and plasmid construction details are described in Supplementary Table 2 and 3.

### Three-plasmid dual-color recombination reporter assay

To engineer the reporter system, PyMOL was used to identify flexible loop regions within the sfGFP structure. The donor sequence was inserted into these regions to generate three distinct structural variants. Each donor-inserted sfGFP variant was cloned into a kanamycin-resistant expression vector carrying a pBBR1 origin (*oriV*), and the constructs were screened to identify mutants that retained normal sfGFP fluorescence despite the insertion (Fig. S4). Functional validation in *P. putida* using a dual-plasmid system confirmed that sfGFP fluorescence was maintained in the basal state but successfully abolished upon recombinase-mediated donor excision (Fig. S5). Based on these results, the optimal sfGFP variant was cloned into a low-copy-number replicating plasmid to serve as the donor vector (pDonor). Concurrently, the target plasmid (pTarget) was constructed by placing the target sequence immediately upstream of the start codon of a promoterless mCherry gene, flanked by two transcriptional terminators, on a separate replicating vector. The expression plasmid (pExp) was designed to co-express the bRNA under a P*_trc_* promoter and the ISPpu10 recombinase under an IPTG-inducible P*_lac_* promoter. The relevant sequences are detailed in Supplementary Table 1.

To perform the assay, all three replicating plasmids, pTarget, pDonor, and pExp were co-electroporated into *E. coli* TOP10. Control strains were established by transforming either the target and donor plasmids together, or the expression plasmid alone. Successful transformation across all strains was verified by colony PCR. The experimental group, control groups, and an untransformed background strain were inoculated into liquid LB medium and cultured overnight (∼12 h) at 37℃ and 220 rpm. Cultures were then subcultured to an initial OD_600_ of 0.2 into fresh LB media containing appropriate antibiotic selections: no antibiotics (background); 100 μg mL^-1^ spectinomycin (expression control); 50 μg mL^-1^ chloramphenicol + 50 μg mL^-1^ kanamycin (target/donor control); 50 μg mL^-1^ chloramphenicol + 50 μg mL^-1^ kanamycin + 100 μg mL^-1^ spectinomycin (experimental group). sfGFP and mCherry fluorescence intensities were measured at 12 h and 24 h as previously described. Insertion efficiency was calculated as the ratio of the increase in mCherry fluorescence in experimental relative to control groups, to the decrease in sfGFP fluorescence over the first 12 h. Finally, to confirm recombination at the molecular level, plasmids were extracted from the remaining cultures and analyzed by PCR using primers spanning the predicted insertion junctions, followed by agarose gel electrophoresis (Fig. S14–15). Primers and plasmid construction details are described in Supplementary Table 2 and 3.

### Prediction of conserved target secondary structures

To identify conserved secondary structures within IS110 insertion sites, LF sequences of IS110 elements were retrieved from the ISfinder database.^16^ The secondary structures and MFE values of these candidate sequences were computed using the ViennaRNA Package (v2.5.0) under physiological conditions (37℃). The resulting structures were computationally filtered to identify stable stem-loops containing a maximum of 1-bp mismatch within the stem region.

### Genome-wide motif mining and spatial analysis

To systematically identify ISPpu10 homologs, a comprehensive dataset comprising 12,096 *Pseudomonas* genomes was curated and phylogenetically clustered at the species level. Intraspecies comparative genomics was performed within these clusters to pinpoint dynamic, non-conserved sequence insertions. Putative insertions were subsequently validated as true ISPpu10 homologs via BLAST searches against a curated reference library. Following validation, the insertion sequences alongside their adjacent left and right flanking regions (LF and RF) were extracted. Multiple sequence alignments of these flanking regions enabled the precise demarcation of transposon insertion boundaries across the dataset, from which a highly conserved target consensus sequence (5 ′ -TTCGCGGGTAAACCCGCTCCTAC-3′) was derived. To investigate the genome-wide spatial organization of these potential target sites, custom Python scripts were utilized to scan the *Pseudomonas putida* KT2440 reference genome, applying the consensus pattern as a query motif under a strict zero-mismatch threshold. Subsequently, a topology-based analysis was conducted focusing on motif orientation and inter-motif spacing. Adjacent motifs were defined as a putative functional pair if they exhibited opposite orientations and were situated within a spatial constraint of 200 base pairs. The genomic distances between these paired motifs were calculated, and their chromosomal contexts (intragenic versus intergenic) were annotated relative to the reference sequence. Finally, statistical analysis was performed to evaluate the preferential enrichment of these functional pairs within intergenic regions compared to coding sequences.

### DBL mutagenesis and colony PCR verification

The donor reprogramming assay was performed in *E. coli* to validate the insertion of the donor plasmid into the target plasmid. The donor plasmid contains an *oriT*, an R6K origin, a kanamycin resistance marker, an sfGFP reporter gene, and the donor sequence. Reliance on the R6K origin renders it a suicide plasmid in *E. coli* TOP10 while allowing stable replication in *E. coli* WM3064, a DAP auxotroph engineered to express the Pir protein.

For donor reprogramming, paired mutations were first introduced at the outermost nucleotide positions of the donor sequence (LD1 and RD5), following the principle of altering base-pairing compatibility. Four variant pairs were designed. Subsequently, the remaining single position, RD1, was mutated to each of the three alternative nucleotides. The target plasmid and the expression plasmid were identical to those used in the three-plasmid system described above. To initiate the experiment, the suicide donor plasmid was electroporated into *E. coli* WM3064, and successful transformation was confirmed by colony PCR. Concurrently, the target and expression plasmids were electroporated into *E. coli* TOP10 and similarly verified. Once verified, the donor strain *E. coli* WM3064 was cultured in LB medium supplemented with 0.3 mM DAP and 50 µg mL^-1^ kanamycin, while the recipient strain *E. coli* TOP10 was cultured in LB containing 100 µg mL^-1^ spectinomycin and 50 µg mL^-1^ chloramphenicol. Both strains were grown overnight for approximately 12 hours in liquid medium at 37℃ and 220 rpm.

For conjugation, 1.5 mL of each culture was centrifuged at 6,000 rpm for 1 min. The supernatant was discarded, and the pellet was resuspended in 1 mL of PBS. This wash step was repeated twice. Finally, the donor and recipient pellets were each resuspended in 100 µL of PBS, mixed thoroughly, and spotted onto an LB agar plate supplemented with 0.03 mM DAP and 5 µg mL^-1^ kanamycin for a 12-h mating period. After conjugation, cells were scraped from the mating spot, resuspended in 100 µL of PBS containing 10 mM IPTG, and plated onto a selection LB agar plate containing 50 µg mL^-1^ chloramphenicol, 50 µg mL^-1^ kanamycin, and 100 µg mL^-1^ spectinomycin. This plate was incubated at 37℃ for 48 h. Due to the absence of DAP alongside the presence of chloramphenicol and spectinomycin, the donor strain cannot survive. Thus, any colonies appearing on this plate represent recipient strains that have successfully integrated the kanamycin-resistant suicide donor plasmid.

To validate the integration, eight random colonies from the selection plate were selected for colony PCR analysis. Two primer pairs were used simultaneously to verify both junction sites of the plasmid insertion. The PCR products were then subjected to Sanger sequencing to determine the precise sequence at the junctions, confirming correct insertion as the LD-CT-RT and LT-CT-RD. The insertion efficiency was calculated as the percentage of colonies the eight selected colonies that yielded both successful junction PCRs and sequence-confirmed expected junctions, based on three independent experiments. Primers and plasmid construction details are described in Supplementary Table 2 and 3.

### *Trans*-mobilization of split-donor DNA substrates

To construct a non-replicative cargo plasmid, the bRNA and ISPpu10 coding sequences within the LE–ISPpu10–RE architecture were replaced with an expression cassette comprising a kanamycin resistance gene and a sfGFP reporter gene. This cassette was subsequently assembled with an R6K replication origin to generate the final non-replicative cargo plasmid. The expression plasmid encoding bRNA and ISPpu10 was first electroporated into *P. putida*, and successful transformation was verified by colony PCR. Verified strains were rendered competent, and 60 µL of competent cells were electroporated with the non-replicative cargo plasmid. Following electroporation, cells were immediately resuspended in 950 µL of antibiotic-free LB medium and allowed to recover at 30°C with shaking at 220 rpm for 2 h. After recovery, cells were pelleted by centrifugation at 6,000 rpm for 1 min, the supernatant was discarded, and the pellet was resuspended in 100 µL of PBS containing 10 mM IPTG. The suspension was then spread onto LB agar plates supplemented with 50 µg mL^-1^ kanamycin and 100 µg mL^-1^ spectinomycin, followed by incubation at 30°C for 48 h. Individual colonies grown on selective plates were streaked and subjected to colony PCR using three primer sets: (i) primers flanking LE, (ii) primers flanking RE, and (iii) primers specific to the *kan*^-^sfGFP expression cassette. Successful excision and chromosomal integration of the LE–*kan*–sfGFP–RE element were indicated by the absence of PCR products with both LE- and RE-flanking primers, together with a positive amplicon from the *kan*-sfGFP cassette. Colonies exhibiting this PCR profile were inoculated into LB broth containing 50 µg mL^-1^ kanamycin and 100 µg mL^-1^ spectinomycin, cultured overnight, and subjected to genomic DNA extraction. Whole-genome sequencing was performed using third-generation sequencing platforms, and the resulting reads were analyzed to confirm precise integration of the LE–*kan*–sfGFP–RE element and to verify that insertion occurred at the predicted wild-type ISPpu10 target locus. Primers and plasmid construction details are described in Supplementary Table 2 and 3.

### Identification and targeting of endogenous *E. coli* genomic loci

To identify specific insertion sites for genomic reprogramming, we screened the *E. coli* MG1655 reference genome (GenBank Accession: NC_000913.3) using a custom Python script based on established recognition rules. The algorithm scanned the entire genome for the target motif (N)*6CTCC(C/T)AC and mapped identified sequences to their genomic coordinates to distinguish between coding sequences (CDS) and intergenic regions. Sites located in intergenic regions were classified as selectable candidates to prevent interference with gene expression; conversely, for sites located within CDS, we evaluated gene essentiality by cross-referencing with the profiling data from a previous study,^26^ retaining only those targets positioned within non-essential editable genes while excluding those in essential genes to ensure cellular viability.

### Molecular docking and dynamics simulations

In the absence of a resolved ISPpu10 crystal structure, the AlphaFold3 web server was first employed to predict the structures of the ISPpu10 dimer-bRNA complex and ISPpu10 dimer-LTG/RTG-LT/RT ternary complex.^27^ The ISPpu10 dimer-bRNA-target DNA assembly were then modeled based on the structural architecture of IS621 to ensure structural accuracy. The stem-loop-free and poly(A) mutant models were generated by removing the stem-loop and introducing mutations at the corresponding positions of the full-length target DNA structure. CHARMM-GUI was used to prepare all of the computational assays containing ISPpu10 complexes.^28^ Each molecular assembly was immersed in a box of TIP3P water with a 150 mM NaCl concentration.

All assemblies were first minimized and then equilibrated under the isothermal-isobaric ensemble (300 K and 1.0 atm) using NAMD 3.0 software with the CHARMM36 force field and the TIP3P model.^29–33^ The temperature and pressure were controlled by Langevin dynamics and the Langevin piston method.^34,35^ Given the considerable size of the simulated systems, the time step was set to 4 fs with hydrogen mass repartitioning.^36^ The total simulation time amounted to about 500 ns for each molecular assembly. A 12 Å cut-off was used to truncate short-range electrostatic and van der Waals interactions. The Particle Mesh Ewald was applied to calculate the long-range electrostatic interaction.^37^ The SETTLE and the SHAKE/RATTLE algorithms were employed to constrain in water and other molecules, respectively, covalent bonds involving hydrogen atoms to their equilibrium length.^38–40^ The r-RESPA multiple-time-step algorithm was used to integrate the equations of motion with a time step of 4 and 8 fs for short- and long-range interactions, respectively.^41^

### Quantification of integration off-target rates

To quantitatively evaluate the off-target rate, we developed a custom two-round mapping bioinformatics pipeline leveraging in silico pseudo-reference sequences representing expected integration geometries: a donor-only reference (DD_ref) comprising the core donor region (DR) flanked by 150-bp donor sequences, alongside left (TD) and right (DT) hybrid references containing 150-bp target genomic flanks fused to the donor sequences. Raw paired-end sequencing reads were initially aligned to the DD_ref using BWA-MEM^22^ (v0.7.17; parameters -K 10000000 -t 8), followed by strict SAMtools^23^ filtering to retain only high-quality primary alignments (MAPQ ≥ 10) by excluding unmapped, secondary, and supplementary reads. To identify candidate chimeric junction reads indicative of structural variations, CIGAR strings were parsed to extract alignments exhibiting a minimum of 20-bp contiguous donor matches and exactly one soft-clipped end of ≥ 15 bp, further requiring the split-read breakpoint to map strictly within a 10-bp tolerance window of the theoretical integration coordinate (position 151) to eliminate false positives. The soft-clipped sequences containing the flanking context were then subjected to a secondary mapping strategy against the corresponding hybrid references (TD for left junctions; DT for right junctions) for systematic classification. Reads completely mapping to the hybrid references with terminal soft-clips < 15 bp were classified as precise on-target integrations, whereas reads failing to align, exhibiting low mapping quality (MAPQ < 10), or retaining substantial soft-clipping (≥ 15 bp) indicative of divergent flanking sequences, were categorized as off-target integrations. Finally, a 10-bp sequence motif immediately flanking the integration breakpoint was extracted from the soft-clipped region of each read to calculate motif frequencies, and the integration off-target rate for each junction was computed as the proportion of off-target reads relative to the total identified integration reads.

### Absolute quantification of insertion events

To map and quantify donor insertion events, we developed a custom strand-aware pipeline. Raw paired-end reads were processed using Cutadapt^42^ (v5.2) to isolate host-specific sequences. To exclude false positives from unintegrated plasmids, reads containing un-recombined wild-type donor junctions were strictly filtered (28-bp overlap threshold), followed by the trimming of 20-bp transposon anchors (15-bp overlap, 10% error rate). Remaining host sequences shorter than 30 bp were discarded. The purified left- and right-flank reads were independently aligned to sample-specific reference genomes using BWA-MEM^22^ (v0.7.17), and filtered with SAMtools^23^ to retain only high-quality, primary mappings (MAPQ ≥ 20; Flag -F 2308). For absolute quantification, exact insertion boundaries were extracted using a strand-aware coordinate strategy via pysam^43^, defining breakpoints based on read orientation and flank direction. Breakpoints with a minimum read depth of 3 were matched across left and right flanks within a ±5-bp spatial window to account for target site duplications. Finally, the mean depth of matched flanks was normalized to Reads Per Million (RPM) against the total raw reads to determine the absolute insertion frequency. The nucleotide distance for all insertion sites was calculated using the Hamming distance metric.

## DATA AVAILIBILITY

The whole-genome sequencing data that support the findings of this study have been deposited in the NCBI Sequence Read Archive (SRA) under accession code PRJNA1439739. The custom code, algorithms, and computational pipelines used in this study are available at GitHub (https://github.com/fogbrid/ISPpu10) and have been archived at Zenodo under DOI https://doi.org/10.5281/zenodo.20771710 (version ISPpu10_code_V1.0; Creative Commons Attribution 4.0 International license). All additional data supporting the conclusions of this study are available within the manuscript and its Supplementary Information files. Source data are provided with this paper.

## AKNOWLEDGEMENTS

We thank Zhouqing Luo (Xiamen University) for helpful discussions and Tianmin Wang (ShanghaiTech University) for critical reading of the manuscript and valuable suggestions. We also thank Dr. Alissa Bleem and Tracy Hodges (National Laboratory of the Rockies) for their assistance in obtaining Nanopore sequencing data of the evolved isolates. We acknowledge the Core Facility of Biomedical Sciences at Xiamen University for analytical support. This work was supported by the National Natural Science Foundation of China (Grants No. 3250010532; 32571662), the Fujian Provincial Natural Science Foundation of China (Grant No. 2026J009007), and the Xiamen Municipal Bureau of Science and Technology (Grant No. 3502Z202471027). All main figures and supplementary figures 3 and 8–11 were generated using BioRender.com.

## AUTHOR CONTRIBUTIONS

X.M.N., B.T.C., and C.L. conceived the project. X.M.N., B.T.C., H.L., Y.Z., X.C., Z.Z., and C.L. contributed to the experimental design. X.M.N., B.T.C., Y.T.L., T.J.C., Y.Y., Y.H.Z., N.Y., Y.N.X. and F.Z. performed the laboratory experiments. X.M.N. and B.T.C. conducted the bioinformatics analyses. H.L. performed the molecular modeling and simulations. C.L. and Z.Z. supervised the research. X.M.N., B.T.C., H.L., Z.Z., and C.L. wrote the manuscript with input from all other authors.

## SUPPLEMENTARY FIGURES

**Fig. S1.**
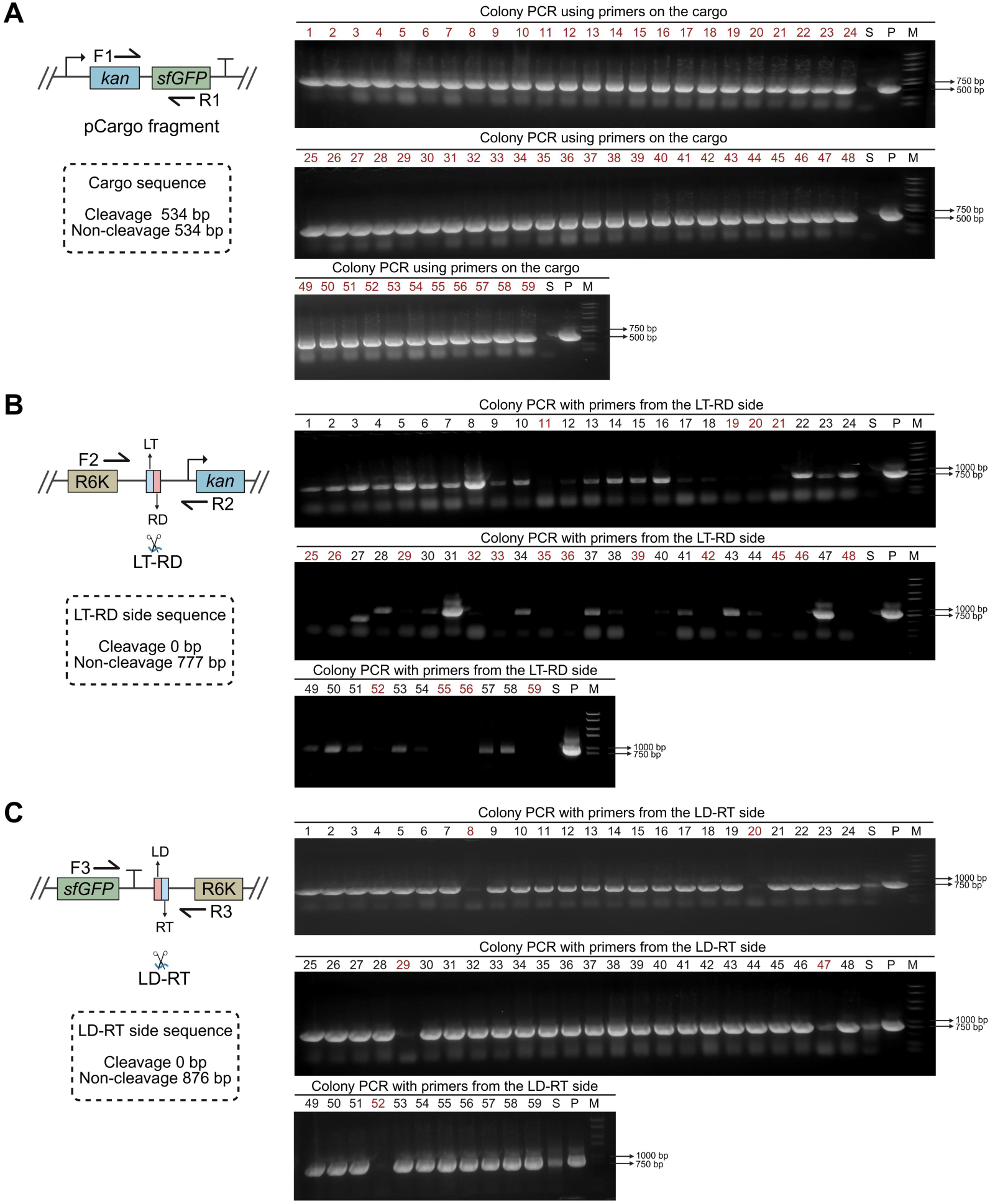
PCR validation of the split-donor capture mechanism. **A.** Colony PCR using primers from the cargo sequence to confirm integration of the cargo into the strain. **B.** Colony PCR using primers from the LT-RD side of the plasmid to verify separation of the cargo sequence from the plasmid vector. If separated, no band will be amplified. **C.** Colony PCR using primers from the LD-RT side of the plasmid to further confirm separation of the cargo sequence from the plasmid vector. If separated, no band will be amplified. The strains (No. 20, 29, and 52) whose colony PCR results with all three primer pairs matched the theoretical expectations were subjected to genomic DNA extraction and long-read sequencing, confirming successful insertion of the cargo into the target genomic locus. P means cargo plasmid, S means plasmid-free colony, M means DNA marker. The numbers marked in red font indicate positive colonies.

**Fig. S2.**
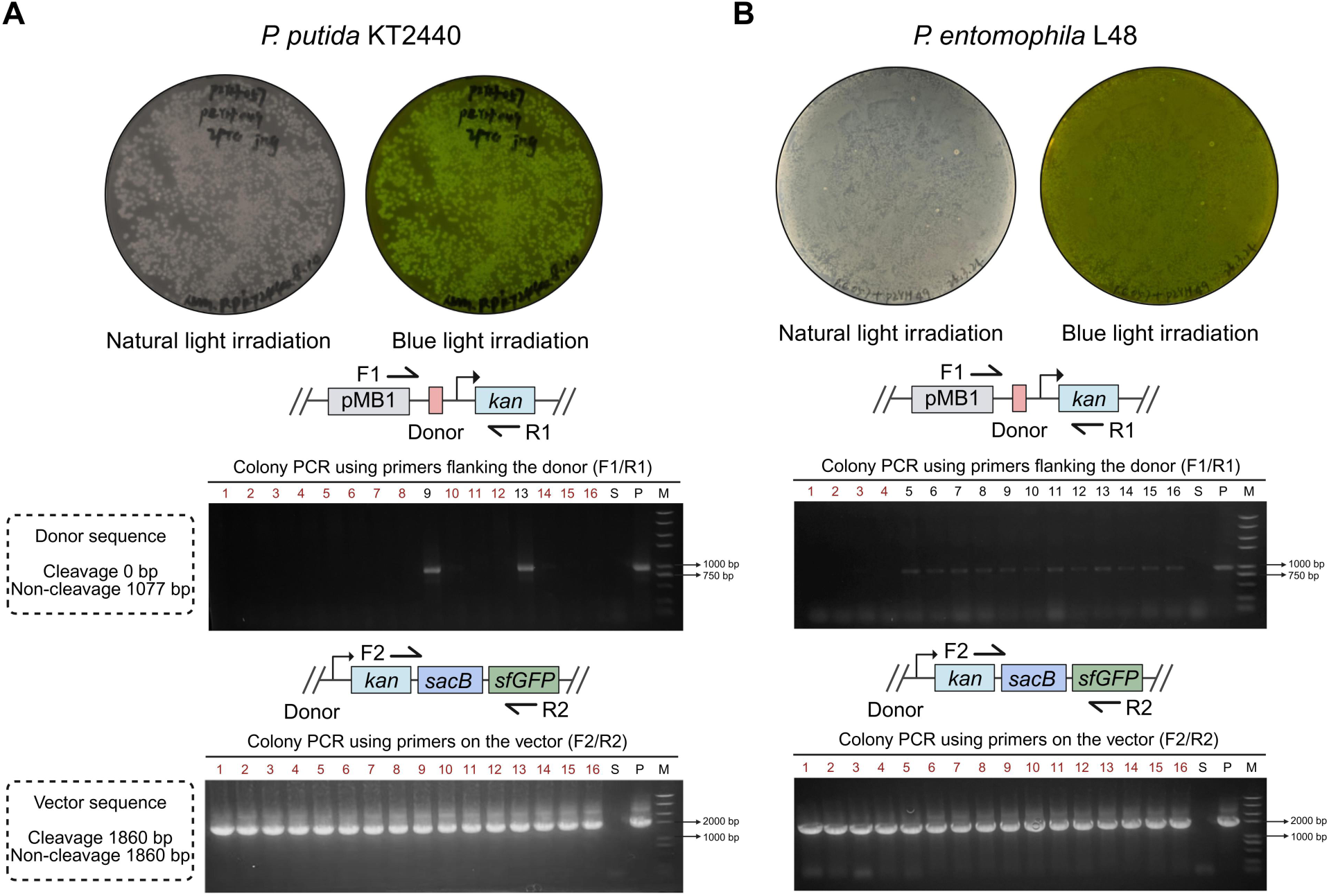
Site-specific genomic integration in *P. putida* and *P. entomophila*. **A.** Plate of single colony grown after integration of the suicide donor plasmid into the target site in the *P. putida* genome, along with colony PCR verification using donor-flanking primers and suicide plasmid vector primers. **B.** Plate of single colony grown after integration of the suicide donor plasmid into the target site in the *P. entomophila* L48 genome, along with colony PCR verification using donor-flanking primers and suicide plasmid vector primers. S means plasmid-free colony, P means the suicide plasmid, M means DNA marker. The numbers marked in red font indicate positive colonies.

**Fig. S3.**
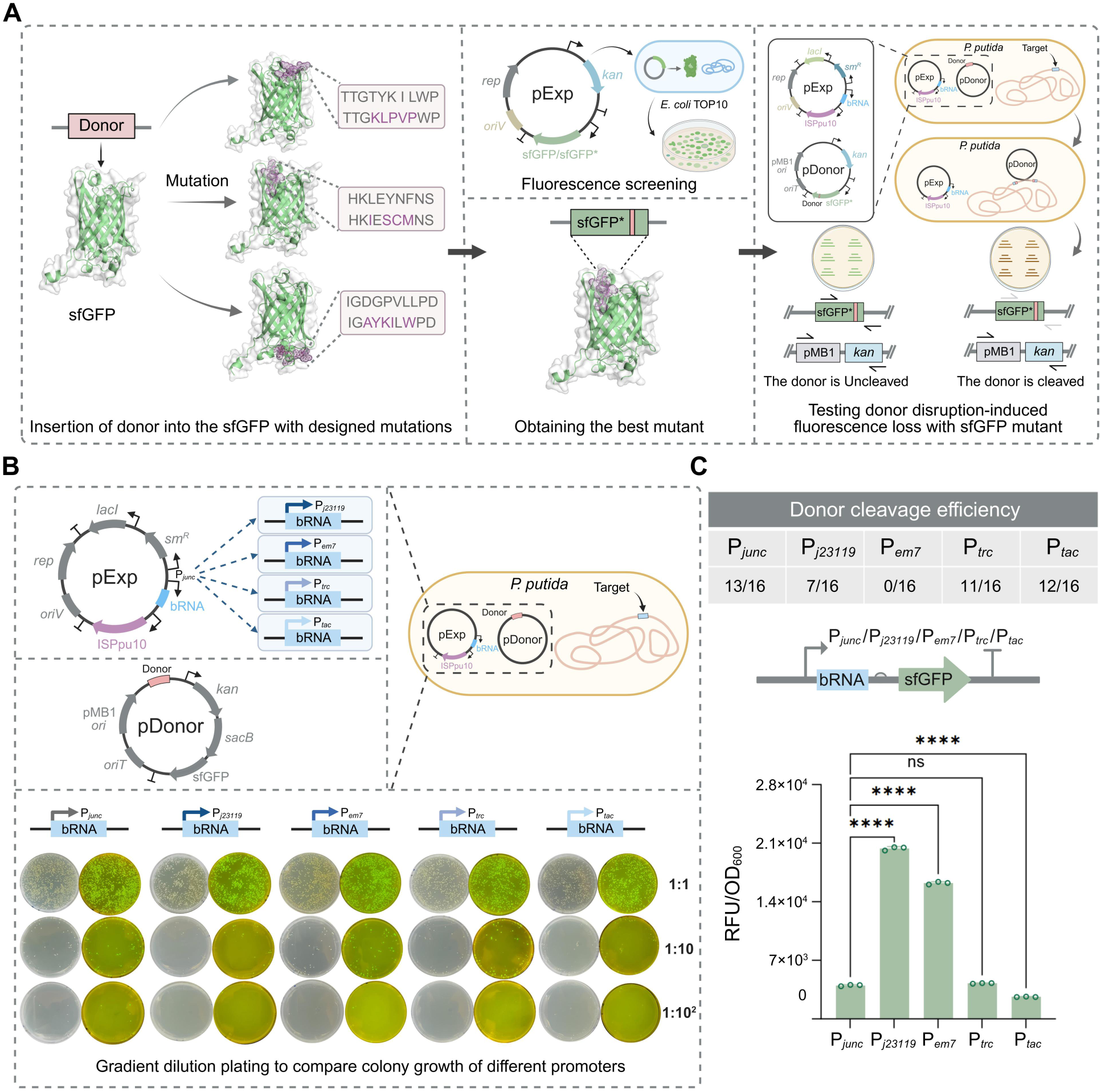
Engineering and optimization of the quantitative three-plasmid reporter system. **A.** Structure-guided engineering of the donor plasmid. The donor DNA sequence was inserted into a permissive loop of the sfGFP coding sequence. Fluorescence screening identified a variant that retains high signal intensity prior to recombination, enabling a loss-of-signal assay for donor excision. **B.** Optimization of the expression plasmid promoter to eliminate inter-plasmid crosstalk. The native P*_junc_* promoter, which inherently contains donor end sequences, mediates unintended plasmid-to-plasmid recombination (false positives). Screening of heterologous promoters (P*_j23119_*, P*_tac_*, P*_EM7_*, P*_trc_*) reveals that while high-strength promoters like P*_EM7_* yield exclusively false-positive clones, P*_trc_* provides an optimal balance. **C.** Quantification of promoter strength and donor cleavage. Comparison of sfGFP fluorescence reveals a strict hierarchy of promoter strength (P*_j23119_* > P*_EM7_* > P*_trc_* ≈ P*_junc_* > P*_tac_*). This confirms that P*_trc_* maintains native-level expression and donor cleavage efficiency (Fig. S6) while abolishing confounding inter-plasmid insertion events.

**Fig. S4.**
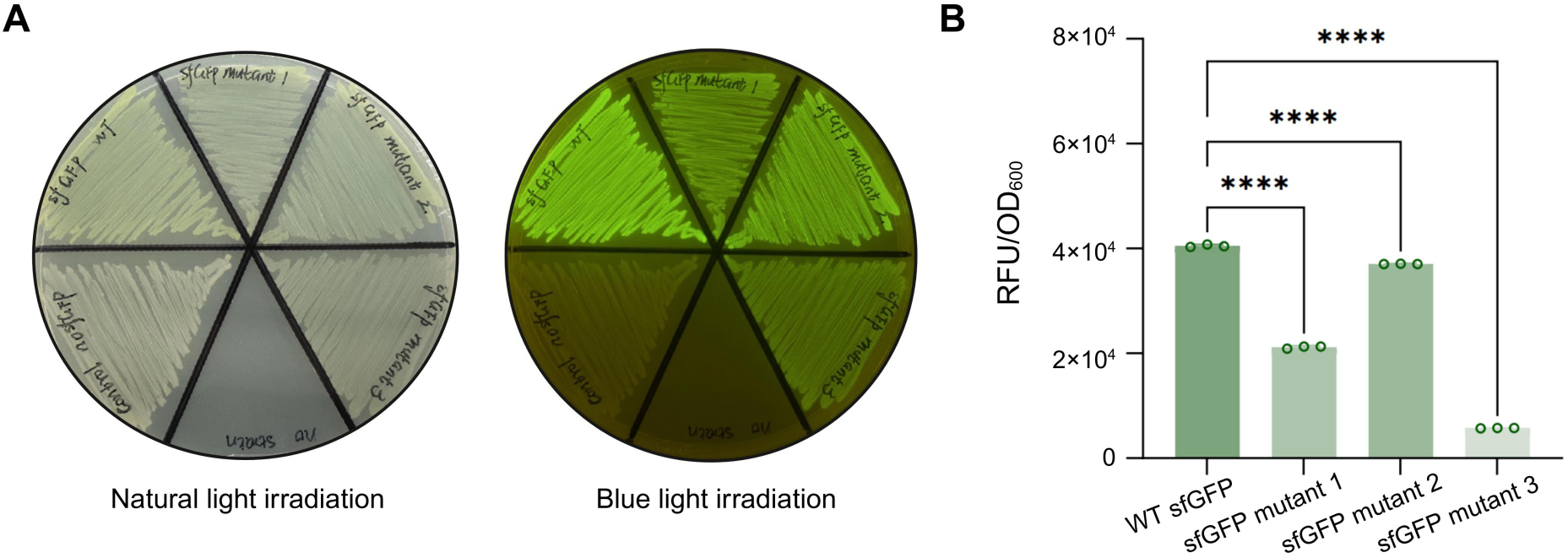
Characterization of fluorescence intensity of sfGFP and its mutants. **A.** Plate schematic showing fluorescence intensity of sfGFP and its mutants under natural light and blue light. **B.** Quantitative analysis of fluorescence intensity of sfGFP and its mutants. Data are mean ± s.d. (n = 3); **** *p* < 0.0001 (two-tailed Student’s *t*-test).

**Fig. S5.**
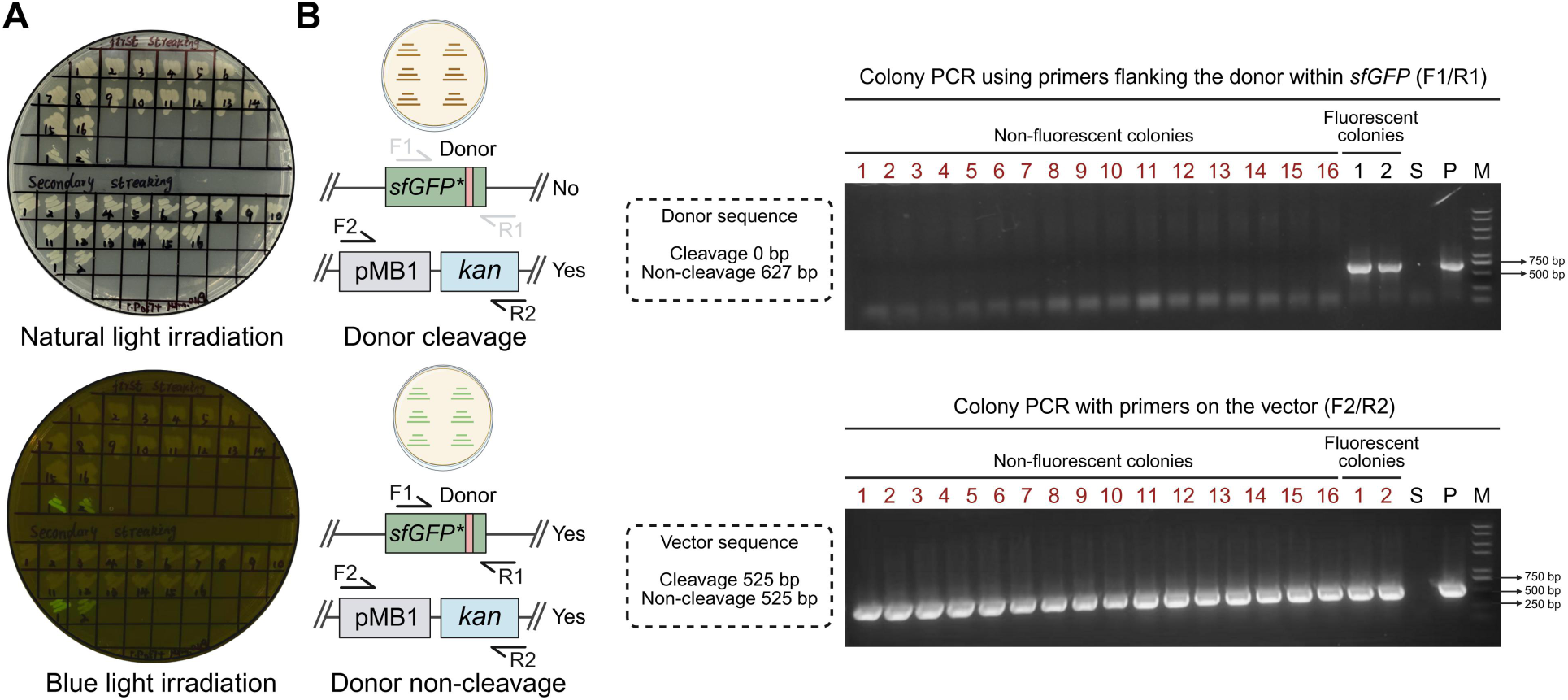
Validation of the reporter activity of sfGFP mutants harboring the donor sequence. **A.** After constructing a suicide plasmid containing an sfGFP mutant with the donor sequence and inserting it into the genome of *P. putida*, the resulting colonies were streaked on plates and their fluorescence intensity was observed under natural light and blue light. **B.** Colony PCR verification using primers flanking the donor and suicide plasmid vector primers. Uncleaved donor colonies exhibit green fluorescence, whereas cleaved donor colonies are non-fluorescent. S means plasmid-free colony, P means the suicide plasmid, M means DNA marker. The numbers marked in red font indicate positive colonies.

**Fig. S6.**
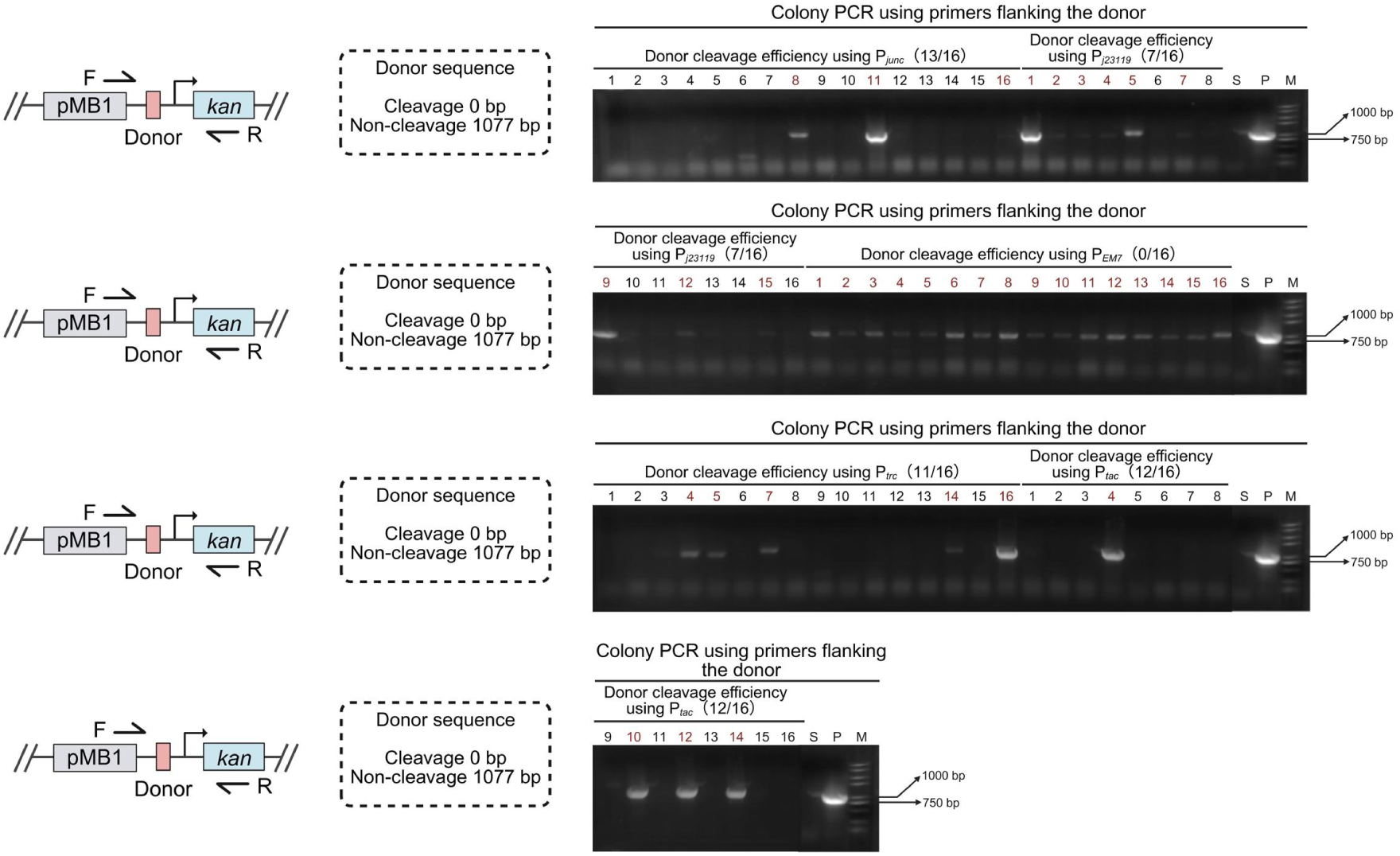
Quantification of promoter strength by colony PCR-based assessment of donor cleavage frequency. Colony PCR-based assessment of promoter strength by quantifying donor cleavage frequency. S means plasmid-free colony, P means the suicide plasmid, M means DNA marker. The red font means the clone numbers in which the donor remains uncleaved.

**Fig. S7.**
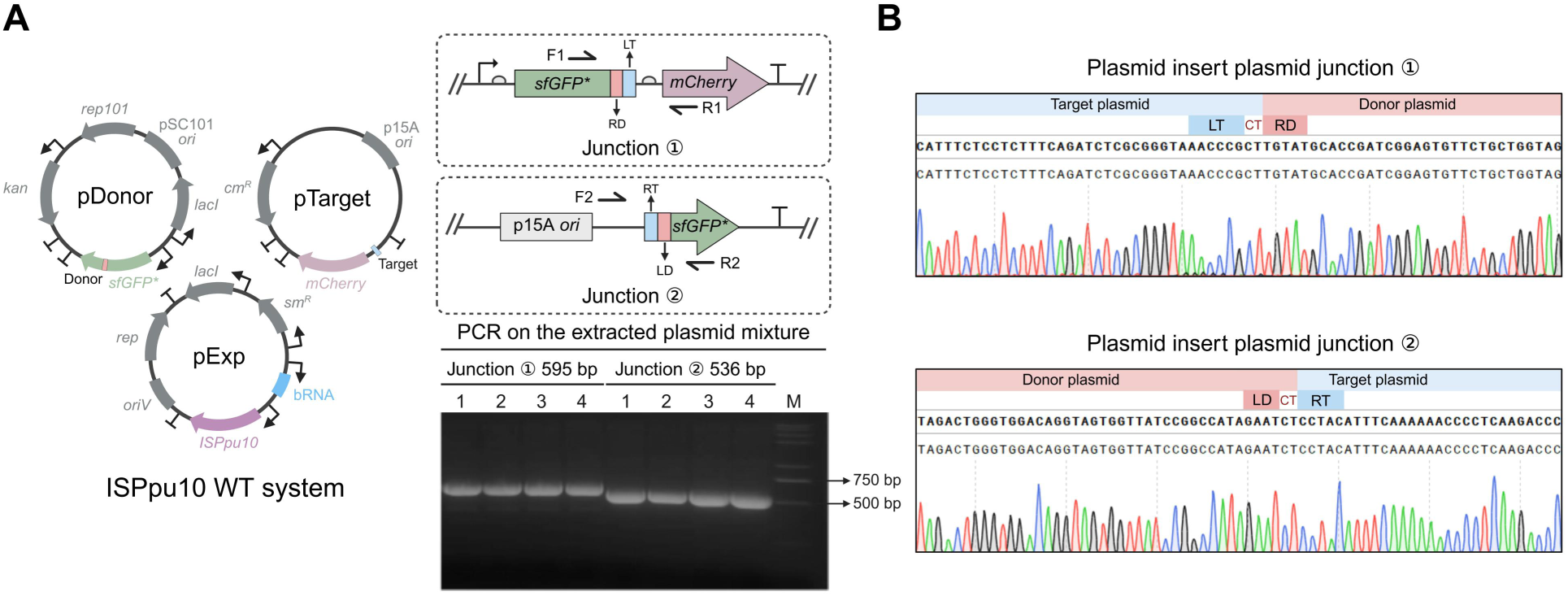
Validation of donor plasmid insertion into target plasmid activity using the three-plasmid WT system in *E. coli* TOP10. **A.** PCR verification of junction ① and junction ② using the plasmid mixture extracted after donor plasmid insertion into the target plasmid in the ISPpu10 WT three-plasmid system. **B.** Sanger sequencing verification of junction ① and junction ②. M means DNA marker.

**Fig. S8.**
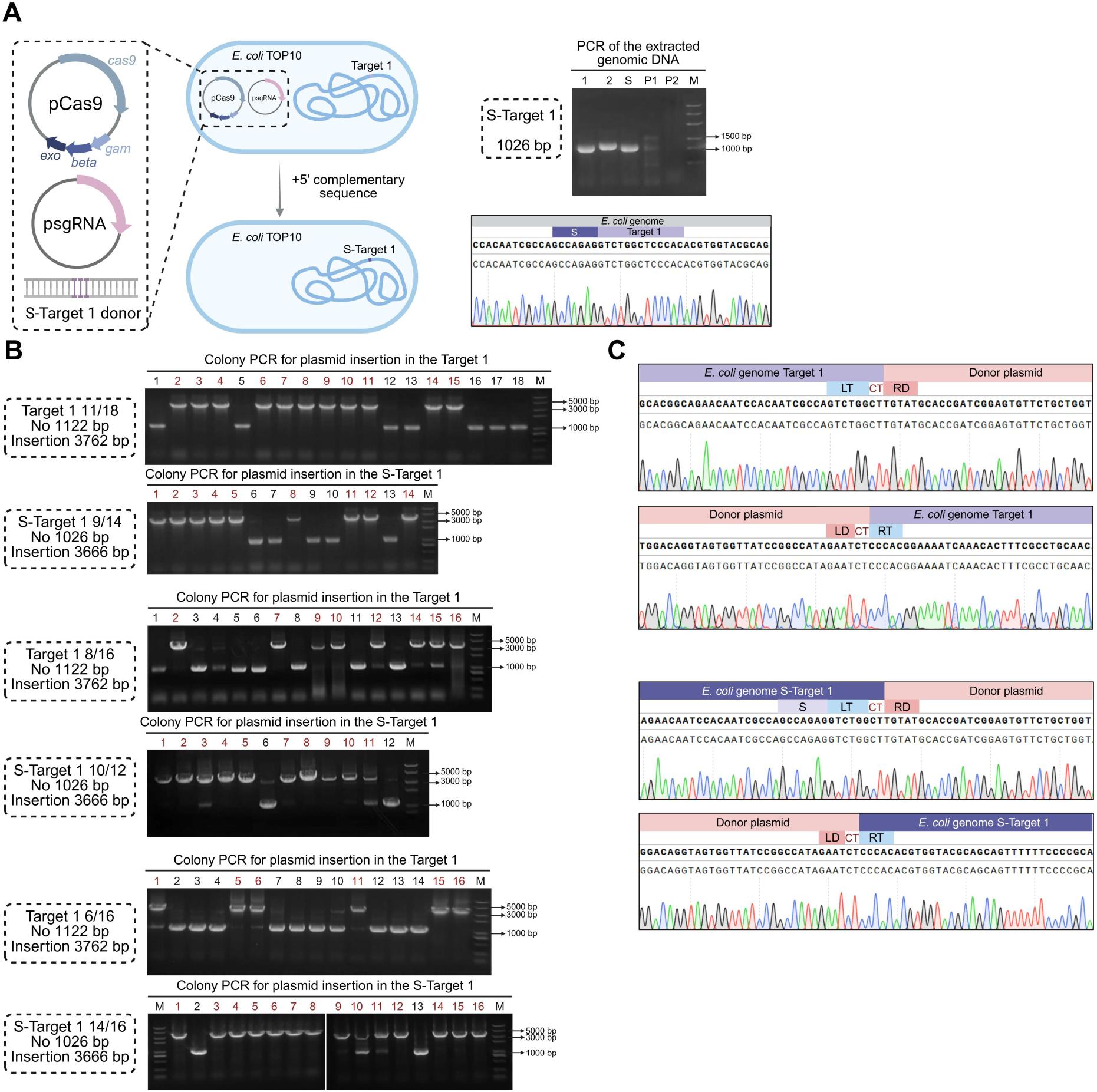
Insertion validation and efficiency of a suicide donor plasmid at reprogrammed loci in the *E. coli* genome. **A.** Construction of the *E. coli* strain with a 5′ complementary sequence added at the reprogrammed Target-1 locus (S-Target 1) in the genome. Successful construction was verified by PCR and Sanger sequencing. P1 means Cas9 plasmid, P2 means sgRNA plasmid, S means plasmid-free colony, M means DNA marker. **B.** Colony PCR validation of colonies resulting from insertion of the suicide donor plasmid into the Target-1 and S-Target 1 loci in the *E. coli* genome, including three independent replicate experiments. M means DNA marker. The numbers marked in red font indicate positive colonies. **C.** Sanger sequencing of PCR fragments from positive colonies at both junctions of the donor inserted into the target site.

**Fig. S9.**
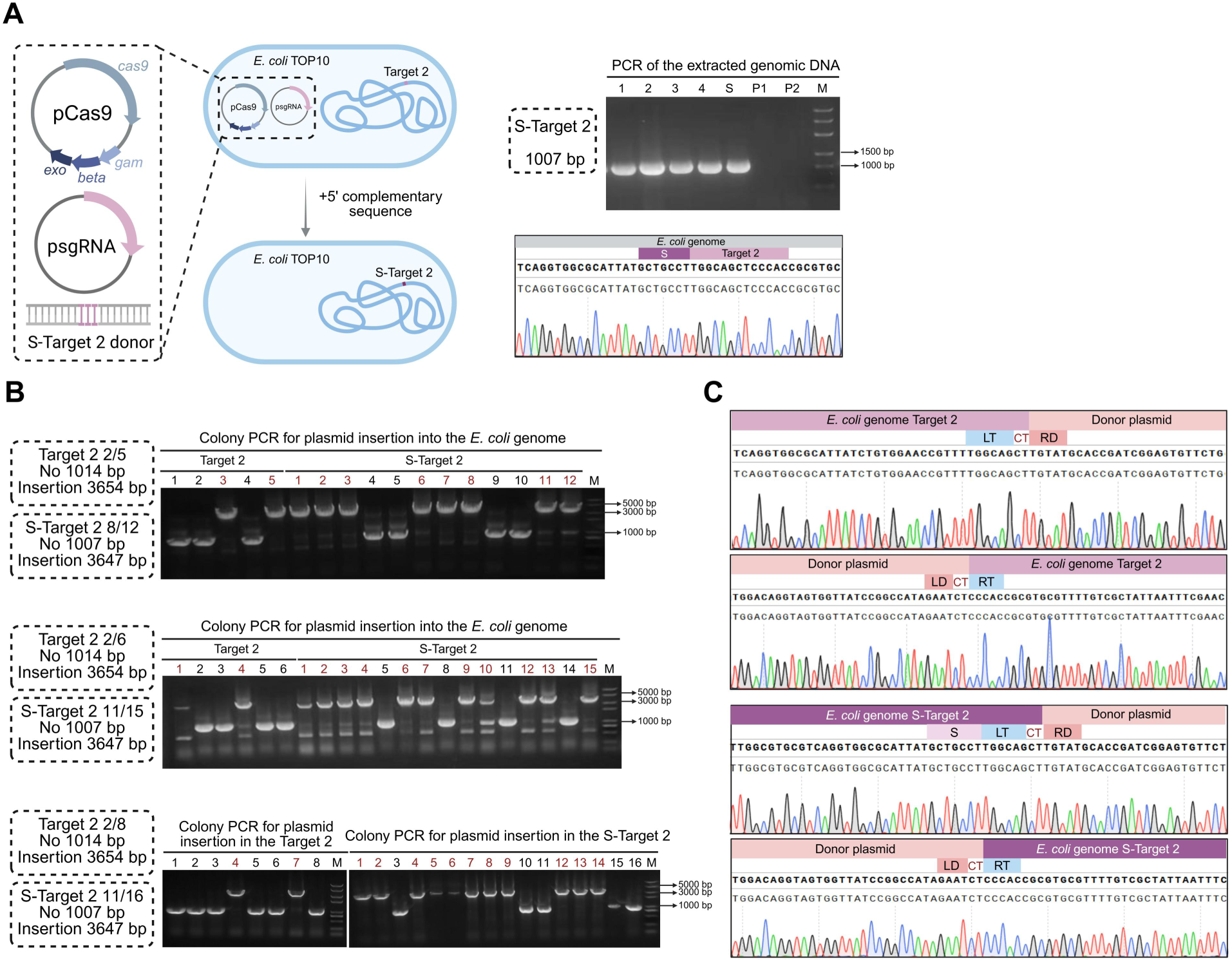
Insertion validation and efficiency of a suicide donor plasmid at reprogrammed loci in the *E. coli* genome. **A.** Construction of the *E. coli* strain with a 5′ complementary sequence added at the reprogrammed Target-2 locus (S-Target 2) in the genome. Successful construction was verified by PCR and Sanger sequencing. P1 means Cas9 plasmid, P2 means sgRNA plasmid, S means plasmid-free colony, M means DNA marker. **B.** Colony PCR validation of colonies resulting from insertion of the suicide donor plasmid into the Target-2 and S-Target 2 loci in the *E. coli* genome, including three independent replicate experiments. M means DNA marker. The numbers marked in red font indicate positive colonies. **C.** Sanger sequencing of PCR fragments from positive colonies at both junctions of the donor inserted into the target site.

**Fig. S10.**
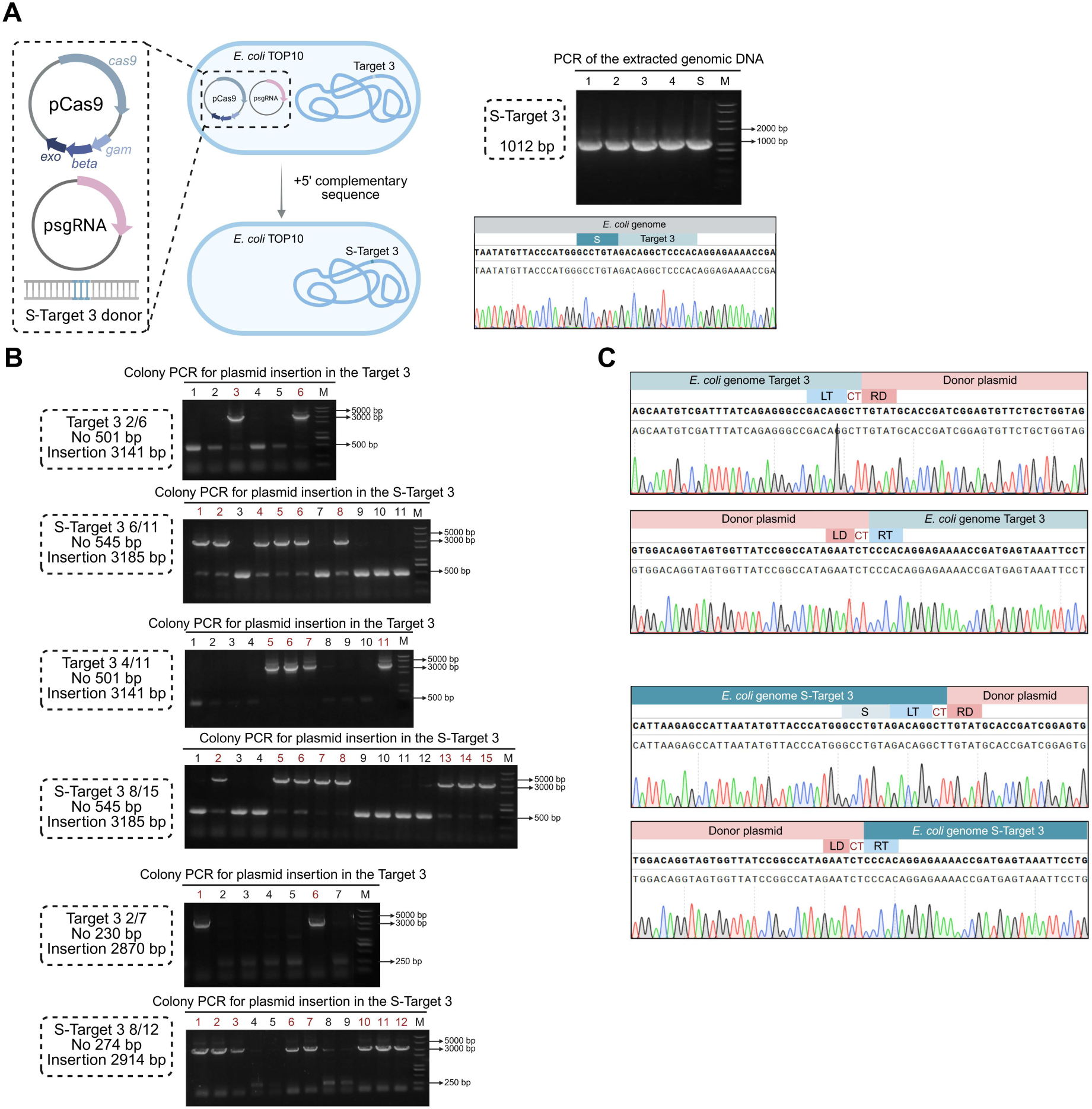
Insertion validation and efficiency of a suicide donor plasmid at reprogrammed loci in the *E. coli* genome. **A.** Construction of the *E. coli* strain with a 5′ complementary sequence added at the reprogrammed Target-3 locus (S-Target 3) in the genome. Successful construction was verified by PCR and Sanger sequencing. P1 means Cas9 plasmid, P2 means sgRNA plasmid, S means plasmid-free colony, M means DNA marker. **B.** Colony PCR validation of colonies resulting from insertion of the suicide donor plasmid into the Target-3 and S-Target 3 loci in the *E. coli* genome, including three independent replicate experiments. M means DNA marker. The numbers marked in red font indicate positive colonies. **C.** Sanger sequencing of PCR fragments from positive colonies at both junctions of the donor inserted into the target site.

**Fig. S11.**
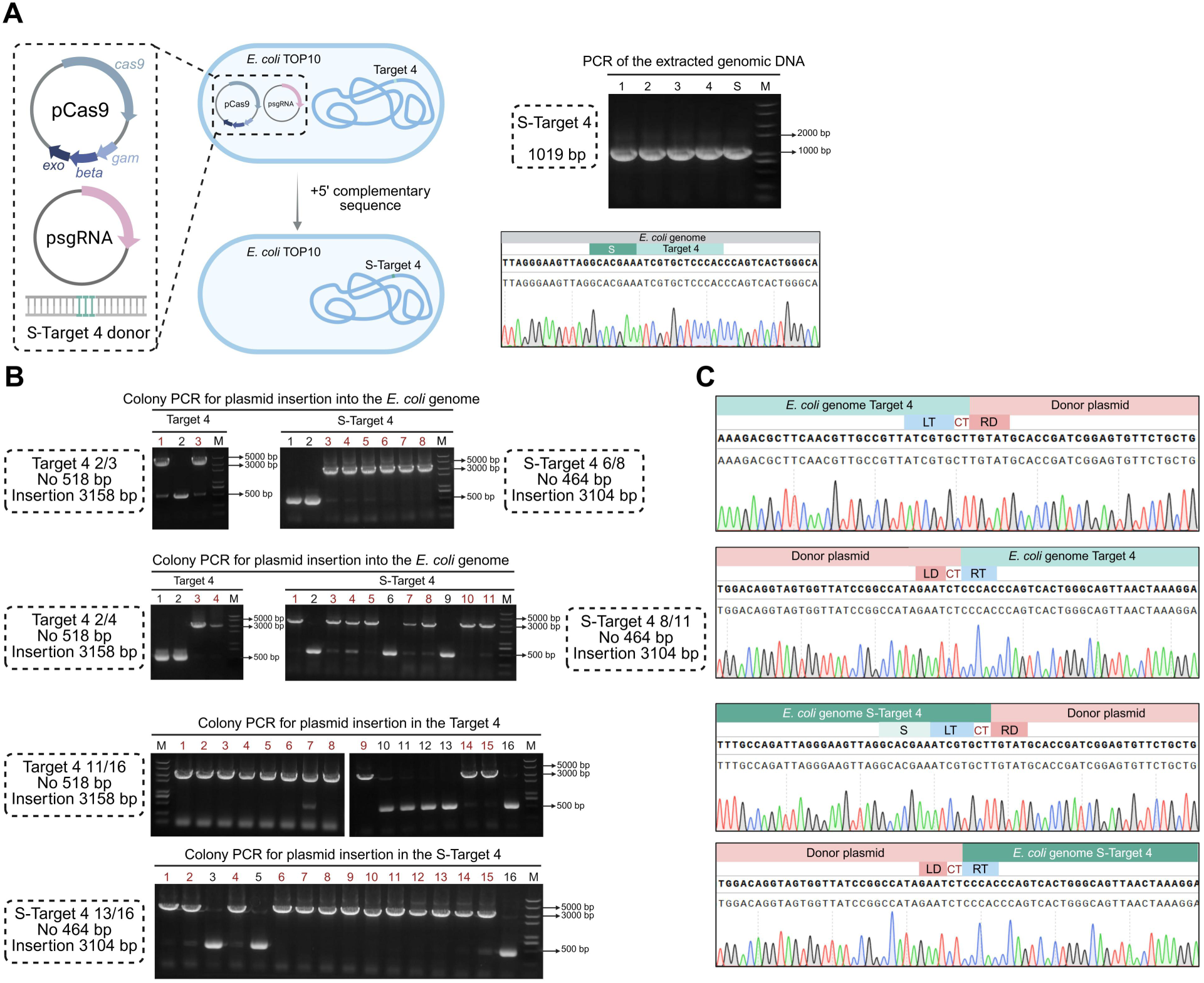
Insertion validation and efficiency of a suicide donor plasmid at reprogrammed loci in the *E. coli* genome. **A.** Construction of the *E. coli* strain with a 5′ complementary sequence added at the reprogrammed Target-4 locus (S-Target 4) in the genome. Successful construction was verified by PCR and Sanger sequencing. P1 means Cas9 plasmid, P2 means sgRNA plasmid, S means plasmid-free colony, M means DNA marker. **B.** Colony PCR validation of colonies resulting from insertion of the suicide donor plasmid into the Target-4 and S-Target 4 loci in the *E. coli* genome, including three independent replicate experiments. M means DNA marker. The numbers marked in red font indicate positive colonies. **C.** Sanger sequencing of PCR fragments from positive colonies at both junctions of the donor inserted into the target site.

**Fig. S12.**
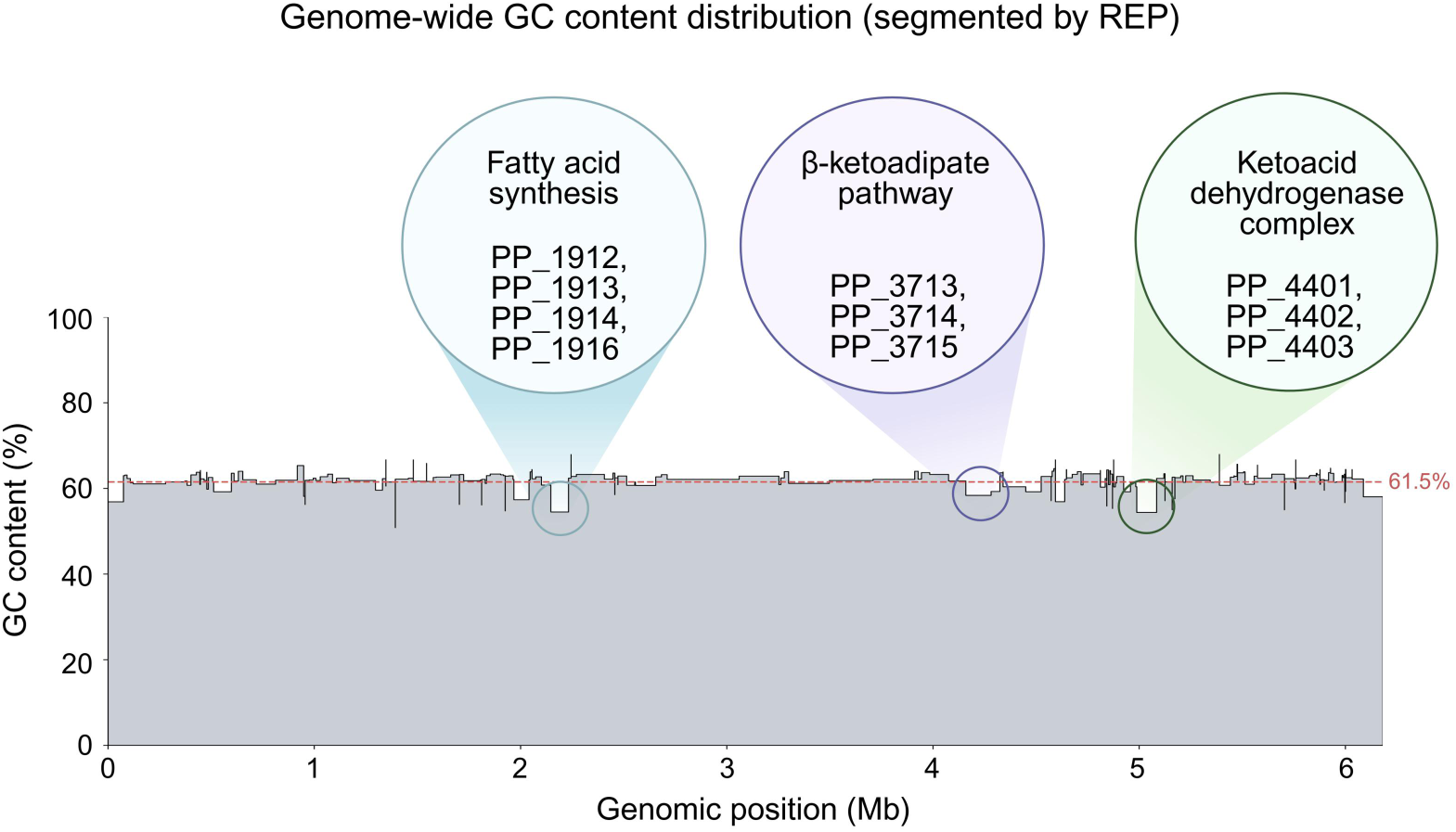
Local GC content across the *P. putida* genome partitioned by REP target sites. The red dashed line indicates the genome-wide average (∼61.5%). Depressions below this baseline mark GC-poor regions. Callouts highlight three such regions encoding complete metabolic pathways: fatty acid biosynthesis (PP_1912–PP_1914 and PP_1916), the β-ketoadipate pathway (PP_3713–PP_3715), and a branched-chain keto acid dehydrogenase complex (PP_4401–PP_4403, bkd genes).

**Fig. S13.**
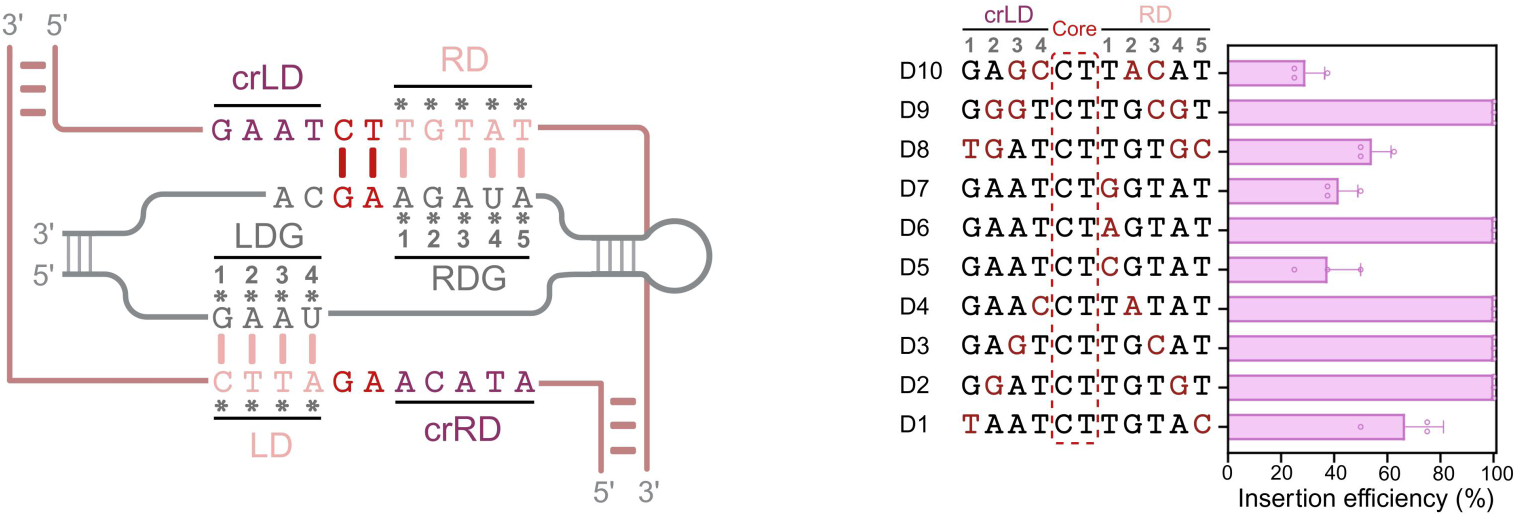
Plasticity of the DBL-dDNA interface. Systematic maximal divergence mutagenesis, involving paired substitutions at the outer donor positions (e.g., LD1 and RD5) and single-nucleotide saturation at RD1, revealed that ISPpu10 mediated plasmid integration across all donor variants tested. Junction PCR and Sanger sequencing confirmed correct insertion junction sequences for the variants, indicating that the DBL–dDNA interface is highly plastic and readily reprogrammable with minimal loss of activity. The positions marked with asterisks indicate the nucleotides to be mutated. Red nucleotides indicate altered positions.

**Fig. S14.**
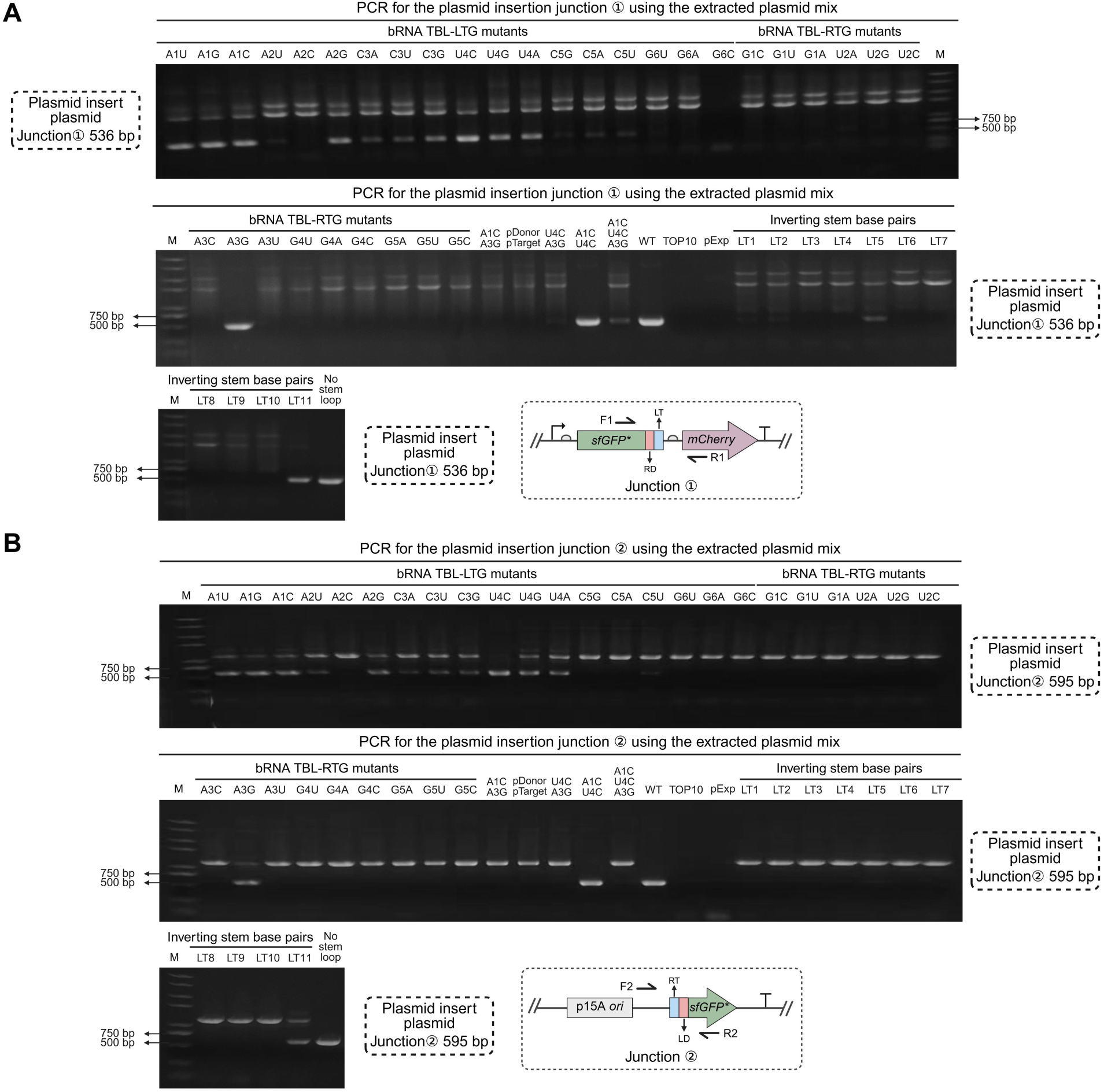
PCR validation of plasmid mixtures extracted from strains carrying the three-plasmid reporter system after donor plasmid insertion into the target plasmid. **A.** PCR verification of junction ①, including the bRNA TBL single-base mutant, the target 5′ stem-loop disruption mutant, and the inverting stem base pairs mutant. **B.** PCR verification of junction ②. pDonor-pTarget refers to the plasmid mixture extracted from control strains containing only the donor plasmid and the target plasmid. pExp refers to the plasmid mixture extracted from control strains containing only the expression plasmid. TOP10 refers to the mixture extracted from plasmid-free control *E. coli* TOP10 strains. WT refers to the plasmid mixture extracted from strains containing the wild-type donor plasmid, target plasmid, and expression plasmid. No stem loop refers to the plasmid mixture extracted from strains containing the donor plasmid, the target plasmid with a disrupted target 5′ stem-loop, and the expression plasmid.

**Fig. S15.**
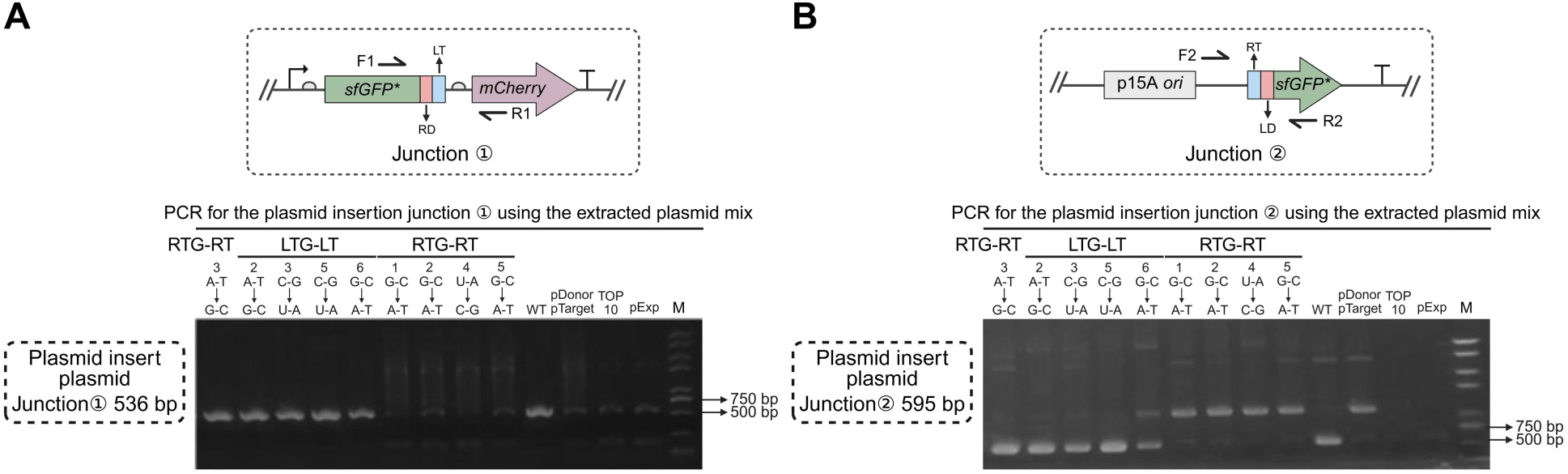
PCR validation of plasmid mixtures extracted from strains carrying the three-plasmid reporter system after donor plasmid insertion into the target plasmid. **A.** PCR validation of junction ① for plasmid insertion using plasmid mixtures from strains with altered bRNA TBL-target base pairing. **B.** PCR validation of junction ② for plasmid insertion using plasmid mixtures from strains with altered bRNA TBL-target base pairing.

